# SIZ1 is a nuclear host target of the nematode effector GpRbp1 from *Globodera pallida* that acts as a negative regulator of basal plant defense to cyst nematodes

**DOI:** 10.1101/725697

**Authors:** Amalia Diaz-Granados, Mark G. Sterken, Jarno Persoon, Hein Overmars, Somnath S. Pokhare, Magdalena J Mazur, Sergio Martin-Ramirez, Martijn Holterman, Eliza C. Martin, Rikus Pomp, Anna Finkers-Tomczak, Jan Roosien, Abdenaser Elashry, Florian Grundler, Andrei J Petrescu, Geert Smant, Aska Goverse

## Abstract

Soil-borne cyst nematodes are obligatory sedentary parasites that cause severe losses to cultivation of major crops such as potato and soybean. Cyst nematodes establish specialised permanent feeding sites within the roots of their host by manipulating plant morphology and physiology through secreted effectors. Here we identified host targets of effector GpRbp-1 and studied their roles in plant-nematode interactions. GpRbp-1 was found to interact in yeast and *in planta* with the potato and Arabidopsis homologues of Siz/PIAS-type E3 SUMO ligase SIZ1. Our results show that a pathogen effector targets the master regulator SIZ1 in plant cells, which has not been demonstrated earlier to our knowledge. The interaction of GpRbp-1 and SIZ1 localizes to the plant nucleus, suggesting that the nuclear functions of SIZ1 as regulator of plant immunity and physiology may be modulated by GpRbp-1. Furthermore, nematode infection assays and transcriptomic profiling indicate that SIZ1 is required for susceptibility to cyst nematodes. So, these data indicate that E3 SUMO ligases may play an important role in plant-nematode interactions. Based on the prediction of SUMO acceptor and interaction sites in GpRbp-1, a model is proposed in which the effector may recruit SIZ1 to be SUMOylated for full functionality in host cells.

**Significance statement:** Here we show that a pathogen effector can target SIZ1, a key protein involved in regulating several aspects of plant biology, most likely to manipulate the SUMOylation of host proteins for successful infection of plants.

## Introduction

Plant parasitic nematodes are small round worms that infect the underground parts of their plant hosts. In agricultural settings, nematode infections cause yearly losses in the order of $157 billion (Abad *et al*., 2008) and it is expected that the rate of nematode infections will increase due to a warmer global climate (Bebber *et al*., 2013). Cyst nematodes are sedentary endoparasites that penetrate and invade the roots of several major food crops from the Solanaceae family as well as cereals, soybean and sugar beet. Cyst nematodes persist in the soil in recalcitrant cysts that contain hundreds of eggs. In the presence of a host plant, infective juveniles (pre-parasitic J2) hatch from the eggs and penetrate the roots by means of an oral needle-like protractible structure, the stylet. Upon penetration, parasitic-J2 migrate intracellularly in the root until they find a suitable cell for establishment of a permanent feeding site and become sedentary. The characteristic permanent feeding site of cyst nematodes, a so called syncytium, is the sole nutrient source that sustains the nematode through three subsequent life stages i.e. J3, J4 and adult (Kyndt *et al*., 2013). Eggs develop inside the body of adult females which eventually die and convert into a hardened cyst.

The interaction with the host plant is vital for the completion of the nematode’s life cycle and it is largely mediated by a suite of effectors secreted by the nematode. However, only a limited set of effectors has been functionally characterised thus far (Juvale & Baum, 2018). Effectors are produced in two oesophageal glands and secreted selectively throughout the different life stages of the nematode to play different roles during the infection process (Hussey, 1989). For example, several plant cell wall-degrading enzymes are secreted by nematodes at the onset of parasitism to modify or degrade plant cell walls, thereby facilitating intracellular migration (Reviewed in (Wieczorek *et al*., 2015)). Also, a number of effectors that mediate reprogramming of the plant cells are required for the initiation, establishment and maintenance of the syncytium (Gheysen & Mitchum, 2011; Mitchum *et al*., 2013; Quentin *et al*., 2013). At the molecular level, nematode secreted effectors function by modifying, competing with, or mimicking the roles of plant structures, genes or proteins. One strategy is the post-translational regulation of host proteins, either directly or indirectly through their interaction with host targets involved in post-translational modification (PTM) (Juvale & Baum, 2018).

Post-translational modifications constitute a powerful tool for functional regulation of proteins in eukaryotic cells (Spoel, 2018; Walsh *et al*., 2005). These regulatory mechanisms rely most often on reversible modifications of peptides and allow a rapid response to variable environmental cues, without requiring gene synthesis (Spoel, 2018). There are different types of post-translational modifications, including the addition of polypeptides onto specific target proteins. The most widely recognised polypeptide addition is ubiquitination, the attachment of several subunits of ubiquitin to target proteins which often function as a molecular marker for protein degradation (reviewed in (Sadanandom *et al*., 2012; Smalle & Vierstra, 2004)). More recently, an additional small peptide was described (SUMO; Small <u>U</u>biquitin-like <u>Mo</u>difier) that bears close structural similarities to ubiquitin and can also be conjugated onto target proteins (Matunis *et al*., 1998). Opposite to ubiquitination, SUMOylation (addition of SUMO) results in variable cellular fates for the target protein. For instance, SUMOylation can alter the subcellular localization, the enzymatic activity or the protein-interaction properties of a target protein (Kurepa *et al*., 2003; van den Burg *et al*., 2010; Augustine & Vierstra, 2018; Verma *et al*., 2018).

The cellular machinery for SUMOylation is largely conserved among eukaryotes, and in plants it is best characterised by studies in the model plant *Arabidopsis thaliana.* Four SUMO isoforms SUMO 1/2/3/5 from *A. thaliana* are shown to be functional, with SUMO1 and 2 as the prevalent isoforms serving as substrate for SUMOylation (Kurepa *et al*., 2003; van den Burg *et al*., 2010). SUMOylation of target substrates is catalysed by a chain of reactions similar to that of ubiquitination (Kurepa *et al*., 2003; van den Burg *et al*., 2010). First, the precursor of SUMO is matured by Ubiquitin-Like Proteases (ULPs) and it is then activated by heterodimeric E1-activating enzymes composed by subunit SAE2 and either SAE1a or SAE1b subunits. The activation of SUMO results in its attachment to the E2 SUMO conjugating enzyme SCE1, which then catalyses the conjugation of SUMO onto an acceptor lysine commonly within the motif ΨKxE in the target protein (Rodriguez *et al*., 2001). Two E3 ligases, SIZ1 and HYP2 seem to act as enhancers of the activity of the E2 conjugating enzyme (Ishida *et al*., 2012). Finally, SUMOylation can be reversed by an isopeptidase activity of the SUMO-activating ULPs (Yates *et al*., 2016).

A large amount of evidence places SUMOylation at the nexus of plant responses to (a)biotic stress (Elrouby *et al*., 2013; Miller *et al*., 2013). For instance, the abundance of SUMO conjugates increases when plants are subjected to heat shock or chemical exposure, including hydrogen peroxide, copper, and ethanol (Chen *et al*., 2011; Kurepa *et al*., 2003). Additionally, Arabidopsis mutants of the different components of the SUMO machinery often display phenotypes defective in tolerance to abiotic stress or pathogen attack (Ishida *et al*., 2012; Kurepa *et al*., 2003; Lee *et al*., 2006; van den Burg *et al*., 2010). In particular, the knock-out mutant of the SUMO E3 ligase SIZ1 has a strong pleiotropic phenotype, indicating that SIZ1 plays a prominent role as regulator in the response to several different types of environmental stresses (Lee *et al*., 2006). In biotic stress, SIZ1 has been shown to be a negative regulator of salicylic acid-mediated defence, i.e. the *siz1-2* knock-out mutant shows increased resistance to infection by *Pseudomonas syringae* pv. *tomato* (Lee *et al*., 2006). Due to its prominent role as a negative regulator of plant immunity, SIZ1 would be a valuable target for pathogen effectors to modulate plant immunity for successful infection of their host. However, no evidence is provided for this hypothesis yet.

The nematode effector GpRbp-1 belongs to the highly expanded family of SPRYSEC proteins of the potato cyst nematodes *Globodera pallida* and *G. rostochiensis* (Diaz-Granados *et al*., 2016; Jones *et al*., 2009; Mei *et al*., 2015; Rehman *et al*., 2009). SPRYSEC effectors contain an N-terminal signal peptide for secretion and a C-terminal SPRY domain. The N-terminal signal peptide suggests that SPRYSEC effectors are delivered to the plant cell where they can interact with host proteins. The C-terminal domain, in turn, is proposed to act as a binding platform to mediate interaction with plant target proteins (Diaz-Granados *et al*., 2016). GpRbp-1 is predominantly expressed during the early parasitic stages of nematode infection, which suggests that it plays a role in early parasitism during the initiation and/or establishment of syncytia (Blanchard *et al*., 2005). A role of GpRbp-1 in nematode virulence is further supported by signatures of positive selection on GpRpb-1 variants from field populations of *G. pallida* (Carpentier *et al*., 2012). The diversification of this effector family is probably due to specific recognition of certain members by the plant immune system, as shown for the potato immune receptor Gpa2. This receptor recognises specific variants of GpRbp-1 and confers resistance to particular populations of *G. pallida* in the field harbouring the corresponding effector variant (Sacco *et al*., 2009).

To elucidate the role of GpRbp-1 in virulence of *G. pallida,* we aimed to characterize its molecular targets in cells of host plants. We used a combination of protein affinity assays to show that the nematode effector GpRbp-1 interacts specifically in yeast and *in planta* with the SP-RING finger domain of a potato Siz1/PIAS SUMO E3 ligase (StSIZ1). Furthermore, we could demonstrate that this interaction occurs in the nucleus of the plant cell. Similarly, GpRBP-1 was able to interact with AtSIZ1, which prompted us to test the role of SIZ1 in cyst nematode infection by using the *Arabidopsis* mutant *siz1-2*. Infection of *in vitro* grown plant resulted in fewer adult nematodes developing on the roots, consistent with the role of SIZ1 as a negative regulator of basal plant defence to biotrophic pathogens (Lee *et al*., 2006). Additional evidence was obtained by a comparative RNAseq analysis, which shows that the reduction of nematode susceptibility in *siz1-2* plants is likely due to the activation of defence-related pathways by the *siz1-2* mutation. So, here we show that an effector from a plant pathogen targets the master regulator SIZ1 to promote disease. Moreover, this study provides evidence for a functional role of SIZ1-mediated SUMOylation in nematode parasitism of plant roots. From our data, a picture emerges in which cyst nematodes target SIZ1 in the nucleus to modulate their host through post-translation modifications. To conclude, possible implications on the modulation of SUMOylation (or SIZ1) in plant cells by cyst nematodes are also discussed.

## Results

### GpRbp-1 interacts in yeast with a fragment of SUMO E3 ligase SIZ1 from potato

To find plant interactors of Gp-Rbp-1 we performed a yeast two-hybrid screen of a cDNA library obtained from potato (SH) roots infected with the potato cyst nematode species *G. pallida.* We screened a library of 3,85×10^6^ clones using a variant of GbRbp-1 from field population Rookmaker (GpRbp-1_Rook-1) as bait. Five yeast clones containing cDNA sequences of 858 – 976bp with identities ranging from 97.5 to 100% were found to interact with bait protein GpRbp-1. To identify the candidate plant target that these clones correspond to, we compared the sequences of all fragments against the UniProtKB/Swiss-Prot nonredundant database using the BLASTX algorithm. All clones showed the highest sequence similarity to *Arabidopsis thaliana* E3-SUMO ligase SIZ1 (e-values 2.11×10^−42^ to 9.6×10^116^) (**Suppl. Table 1**). Among the five yeast clones, there were two pairs with 100% identical sequences within each pair (StSIZfrag10 and StSIZ1frag14; StSIZ1frag49 and StSIZ1frag83). One additional clone contained a fragment that was 87% the length of the fragment contained in the identical clones (StSIZ1frag06). Sequence alignment showed that the clones localized to the C-terminal half of SIZ1 containing a predicted SP-RING finger domain (**Fig. 1; Suppl. Fig. 1**).

**Figure 1.**
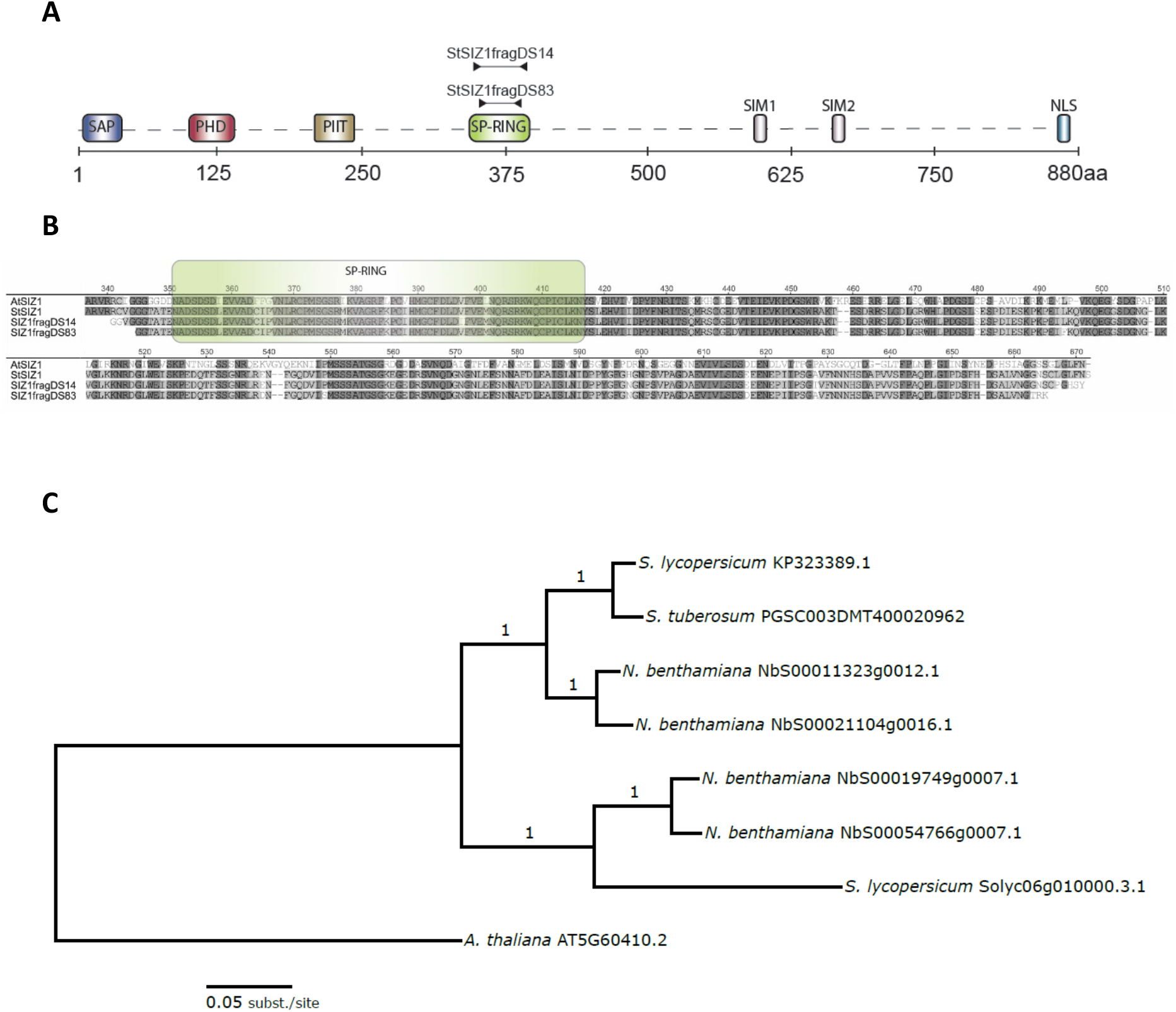
GpRbp-1 interacts with fragments encoding the SP-RING domain of StSIZ1. **A)** Schematic representation (not to scale) of the domain structure of E3-SUMO ligase SIZ1 from *A. thaliana* (AtSIZ1), indicating the SAP, PHD, PIIT, SP-RING, SIM1 and 2 (SUMO interacting) domains and NLS (Miura *et al*., 2005; Rytz *et al*., 2018). Arrows indicate the region where the fragments contained in the yeast clones align to AtSIZ1. **B)** Protein alignment of AtSIZ1(TAIR AT5060410.2), StSIZ1 (XP_006340142.1) and GpRbp-1 interacting fragments (StSIZ1fragl4 and StSIZ1frag83). The alignment was generated using a ClustalW algorithm in Geneious (Geneious version 8.1.9) **C)** (Phylogenetic) tree of StSIZ1 (GenBank XM_015314510.1), predicted NbSIZ1 (Solgenomics Niben101Scf05710g03032.1, Niben101Scf05710g03032.1 and Niben101Scf07109g04008.1), SlSIZ1 (GenBank KP323389.1 and Solyc06g010000.3.1) and AtSIZ1 (TAIR AT5G60410.2). The scale indicates substitutions per site.

In Arabidopsis, the *SIZ1* gene encodes four protein domains, an N-terminal SAP domain (Scaffold attachment factors SAF-A/B, Acinus, PIAS), PHD (Plant Homeodomain), a PIIT (proline-isoleucine-isoleucine-threonine) motif, and a SP-RING (SIZ/PIAS-REALLY INTERESTING NEW GENE). Additionally, two SUMO Interacting Motif (SIM) domains were encoded by *AtSIZ1* (Miura *et al*., 2005) (**Fig. 1**). Finally, AtSIZ1 also contained a nuclear localisation sequence (NLS) in the C-terminal domain of the protein (**Fig. 1**). We compared the coding and peptide sequences of SIZ1 from *Arabidopsis* and potato to investigate their similarity. The full-length coding sequence for *StSIZ1* was obtained from the non-redundant nucleotide database of GenBank (XM_006340080.2). At the nucleotide level, *AtSIZ1* and *StSIZ* shared ~60% identity and at the protein level they shared ~62% identity (**Suppl. Fig. 1**). It should be noted that StSIZ1frag14 and StSIZ1frag83 were 97% identical to StSIZ1, and differences were likely due to the differences in potato genotype used (**Fig. 1B, Suppl. Fig. 1**). Additionally, we investigated the number of copies of *StSIZ1* present in the genome sequence of the doubled monoploid potato genotype DM. To this end, we probed the PGSC *S. tuberosum* group Phureja DM1-3 transcripts v3.4 database from the Potato Genomics resource of Michigan University (Hirsch *et al*., 2014) with a BLASTN algorithm, using AtSIZ as query. Two transcripts (PGSC0003DMT400020963 and PGSC0003DMT400020962; hereafter named 0963 and 0962, respectively) were found corresponding to the same locus in chromosome 11 (PGSC0003DMG400008114), with only transcript PGSC0003DMT400020962 considered to be the representative transcript for the locus (Hirsch *et al*., 2014). Transcript 0963 is 2396bp long, whereas the GenBank StSIZ1 transcript is 3293bp long. An alignment of the protein products for each transcript shows that the peptide encoded by transcript 0962 (StSIZ SpudDB) shares 98% identity to the StSIZ1 GenBank peptide and encompasses the C-terminal half of SIZ1 protein. These results suggest that StSIZ1 is encoded by a single gene residing on chromosome 11 of the DM potato genotype.

### GpRbp-1 interacts with full length StSIZ1 *in planta*

To independently confirm the interaction *in planta,* we used epitope-based co-immunoprecipitation assays. We selected fragments StSIZ1frag14 and StSIZ1frag83 which share a 97% nucleotide identity, differing in ~20 SNPs and 19 nucleotides in length (**Suppl. Table 1**). StSIZ1frag06 shares 100% identity with StSIZ1frag14 and StSIZ1frag83 (**Suppl. Table 1**) and was therefore not used to confirm the interaction *in planta.* GpRbp-1 with an N-terminal Myc-GFP tag (Myc-GFP-Rbp1) was co-expressed with the N-terminally HA-tagged fragments StSIZ1frag14 and StSIZ1frag83 (HA-StSIZ1frag14, HA-StSIZ1frag83) by *Agrobacterium tumefaciens* infiltration in *N. benthamiana* leaves. HA-StSIZ1frag14 and HA-StSIZ1frag83 were specifically co-immunoprecipitated by Myc-GFP-Rbp1 and not by Myc-GFP (negative control) captured by magnetic anti-Myc beads (**Fig. 2A**). Therefore, we concluded that GpRbp-1 interacts *in planta* with two fragments corresponding to a sub-region of the SPRING finger domain of SIZ1 from potato. It is worth noting that after co-immunoprecipitation StSIZ1frag14 and StSIZ1frag83 were detected on western blots as bands migrating approximately 100 KDa higher than the respective bands for the input. This suggests that a complex comprising other peptides may be pulled-down by GpRbp-1.

**Figure 2.**
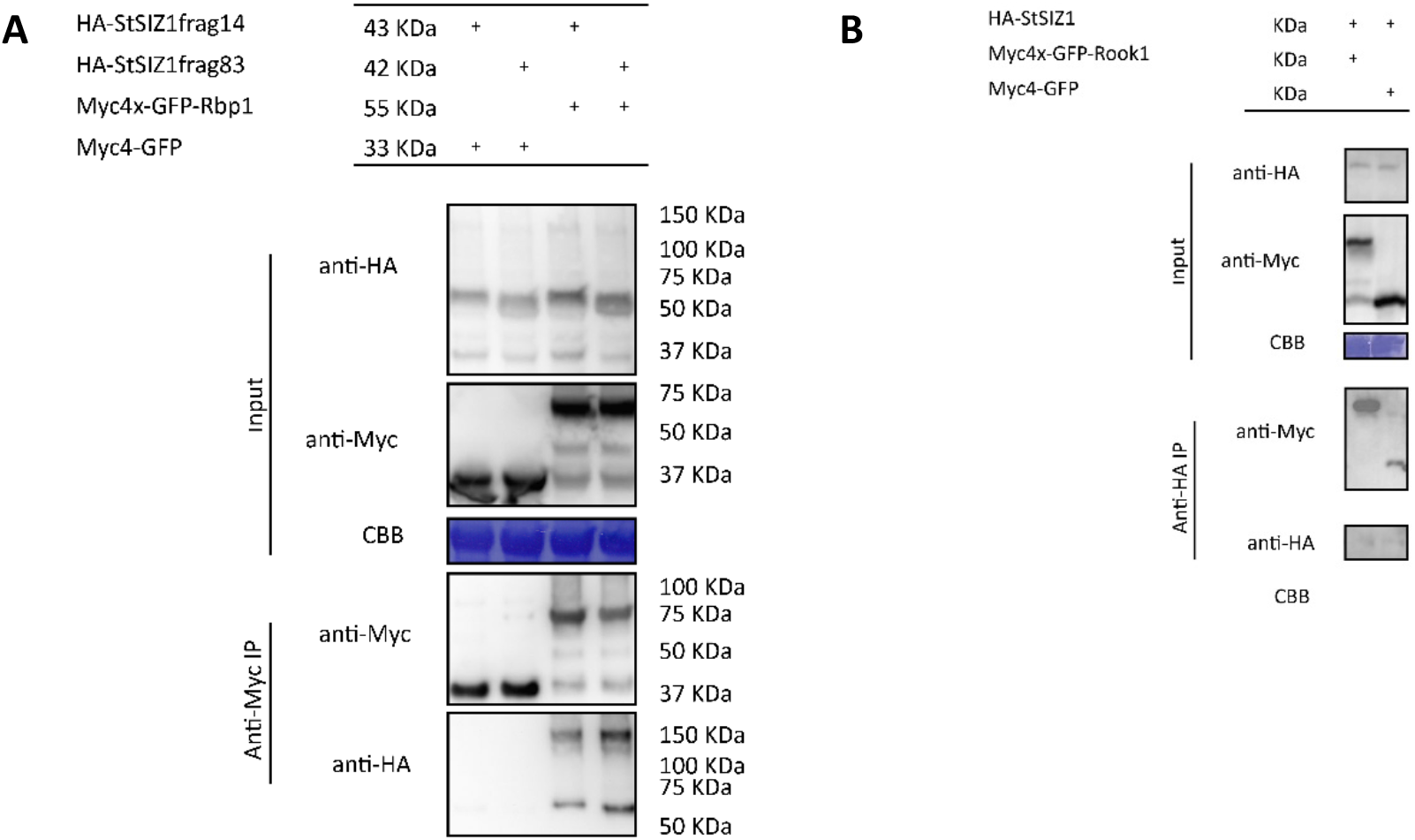
SIZ1 and GpRbp-1 interact *in planta.* **A)** Co-immunoprecipitation of epitope tagged StSIZ1 fragments (HA-StSIZ1frag14/83) and GpRbp-1 or (Myc4-GFP-Rbp1) or empty vector control (Myc4-GFP-EV). Pull-down of Myc-GFP-Rbp1 with anti-myc agarose beads results in a specific co-immunoprecipitation of SIZ1 fragments StSIZ1frag14 and StSIZ1frag83. Co-expressing proteins were extracted from *N. benthamiana* leaves 3 days after agroinfiltration. Results are representative of 3 biological repeats and all co-infiltrations contain the silencing suppressor p19. **B)** Co-immunoprecipitation of full-length HA-tagged StSIZ (HA-StSIZ1) as in **A)**.

To confirm if GpRbp-1 also interacts with full-length SIZ1 from potato, we performed co-immunoprecipitation assays. First, full-length StSIZ1 was obtained by gene synthesis based on the predicted sequence for SIZ1 from potato transcript variant X2 (GenBank code XM_015314510.1 / SpudDb PGSC0003DMT400020962). N-terminally tagged GpRbp-1 (Myc-GFp-Rbp1) was co-expressed with full-length StSIZ1 with an HA-tag (HA-StSIZ1) in *N. benthamiana* leaves by agro-infiltration. Myc-GFP-Rbp1 captured by magnetic anti-Myc beads, co-immunoprecipitated HA-StSIZ1 (**Fig. 2B**). A Myc-GFP (negative control) did not co-immunoprecipitate HA-StSIZ1, indicating a specific interaction of GpRbp-1 with StSIZ1. These results showed that GpRbp-1 was able to interact specifically with StSIZ1 *in planta.*

### StSIZ1 and GpRpb-1 co-localize when expressed *in planta*

A C-terminal nuclear localization signal (NLS) was predicted in StSIZ1. Additionally, SIZ1 from Arabidopsis is exclusively located within the nucleus of cells (Miura, 2005). Therefore, we investigated the localization of StSIZ1 *in planta* by expressing N-terminally GFP-tagged StSIZ1 by transient transformation in *N. benthamiana* leaves. The localization of GFP-StSIZ1 followed a similar nuclear localization as previously reported for AtSIZ1 and the tomato homologue SlSIZ1 (**Fig. 3**) (Lee *et al*., 2006; Zhang *et al*., 2017). Moreover, we observed that the GFP fluorescent signal was uneven throughout the nucleus, with stronger emission in discrete globules within the nucleus, which was also consistent with the localization reported for AtSIZ1. Additionally, StSIZ1 was co-expressed with GpRbp-1 to evaluate their subcellular localization *in vivo* by confocal laser scanning microscopy (CLSM). Co-transformed mCherry-labelled GpRpb-1 (mCh-Rbp1) and GFP-labelled StSIZ1 (GFP-StSIZ1) in *N. benthamiana* leaves were evaluated. When expressed in combination with free GFP, GpRbp-1 was consistently distributed between the nucleus and the cytoplasm as established previously (Jones *et al*., 2009). GFP-StSIZ1 localized to the nucleus with higher expression levels in defined nuclear foci when co-expressed with free mCherry. When co-expressed, GFP-StSIZ1 and mCh-Rbp1 colocalized to the nucleus of transformed cells, suggesting that an interaction occurs in the nucleus. Moreover, we concluded that the subcellular localization of GpRbp-1 or StSIZ1 was not altered upon co-infiltration and apparently, is not affected by their complex formation. The fluorescent tags were fused to the N-terminus of GpRbp-1 and StSIZ1 to simulate as closely as possible the configuration of the proteins in the interaction studies. There, the yeast binding domains or epitope tags were also fused to the N-terminal regions of the CDS of the interactors.

**Figure 3.**
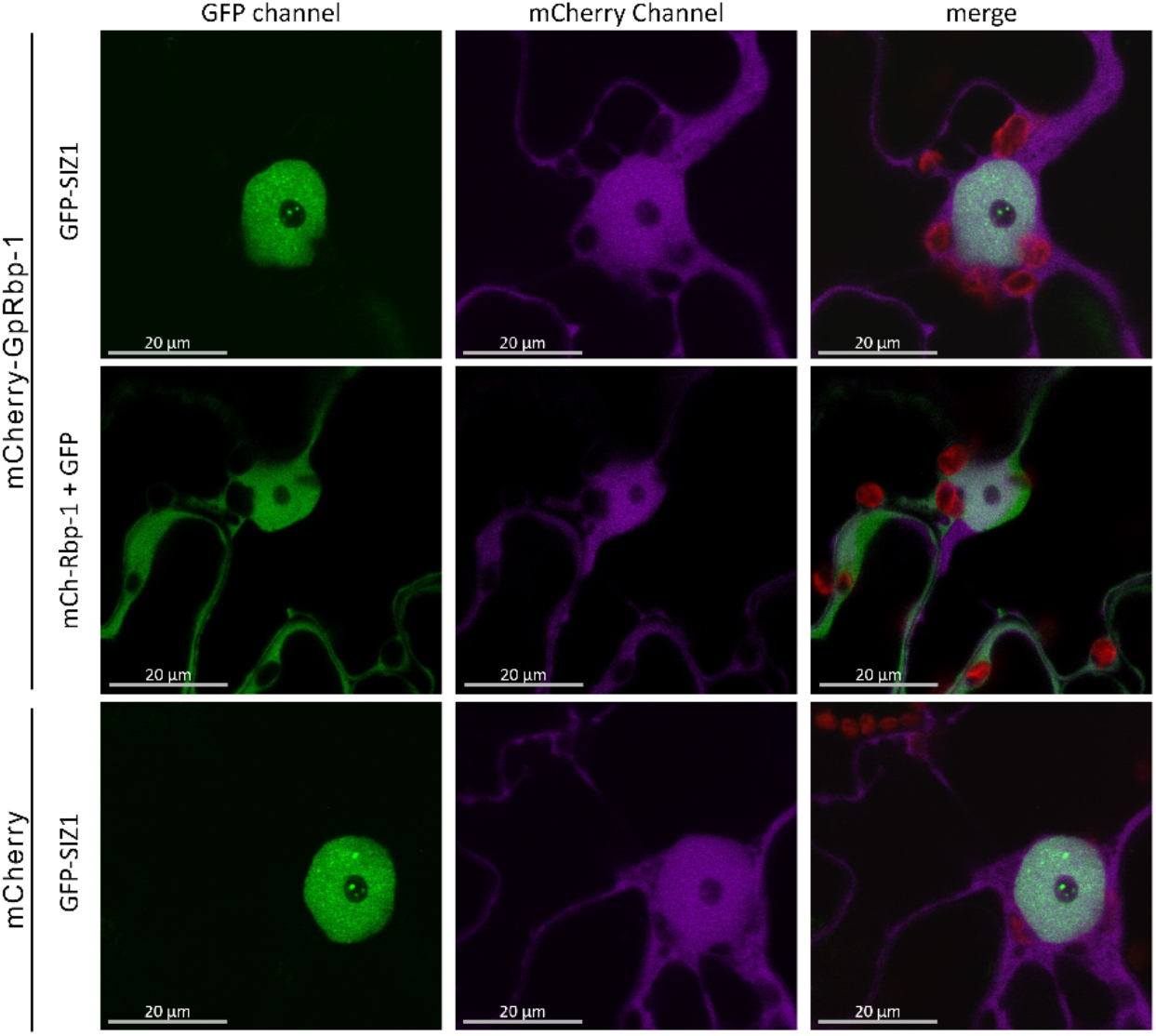
StSIZ1 and GpRpb-1 co-localize to the nucleus of *N. benthamiana* plants. Live imaging of *N. benthamiana* epidermal cells agro-infiltrated with constructs of mCherry tagged GpRbp-1 (mCh-Rbp1) and GFP tagged StSIZ1 (GFP-SIZ1) or GFP and mCh as negative controls. The GFP channel is shown in green and the mCherry in purple. Imaging was performed at 2 dpi, images are representative of 3 biological repeats.

### GpRbp-1 interacts with StSIZ1 and AtSIZ in the plant cell nucleus

To test if GpRbp-1 indeed interacts with StSIZ1 in the nuclear compartment we performed a bimolecular complementation assay (BiFC). For BiFC, the N-terminal half of the super cyan fluorescent protein SCF3A was fused to GpRbp-1 (pN:Rbp1) and the C-terminal half of SCFP3A was fused to StSIZ1 (pC:StSIZ1). The fluorescent fusions were transiently expressed in *N. benthamiana* by agroinfiltration. pN:Rbp1 was co-infiltrated with the viral protein NSs fused to the C-terminal half of protein SCFP3A (pC:NSs) and pC:StSIZ1 was co-infiltrated with β-glucuronidase fused to the N-terminus of SCFP3A (pN:GUS) as negative controls. The characteristic emission of SCFP3A was only reconstituted when pN:Rbp1 and pC:StSIZ1 were co-expressed (**Fig. 4**). There was no reconstitution of the fluorescent signal of CFP when pN:Rbp1 was co-expressed with pC:NSs, neither by the co-expression of pC:StSIZ1 with pN:GUS. These findings confirmed that GpRbp-1 and full-length StSIZ1 interacted specifically *in planta* (**Fig. 2; Suppl. Fig. 1**). Interestingly, the fluorescent signal of CFP was only detected in the nucleus of transformed cells, confirming that GpRpb-1 and StSIZ1 only interact within this cellular compartment. Moreover, the observed granular fluorescent pattern suggests that the interaction between GpRpb-1 and StSIZ1 follows specific substructures within the nuclei.

**Figure 4.**
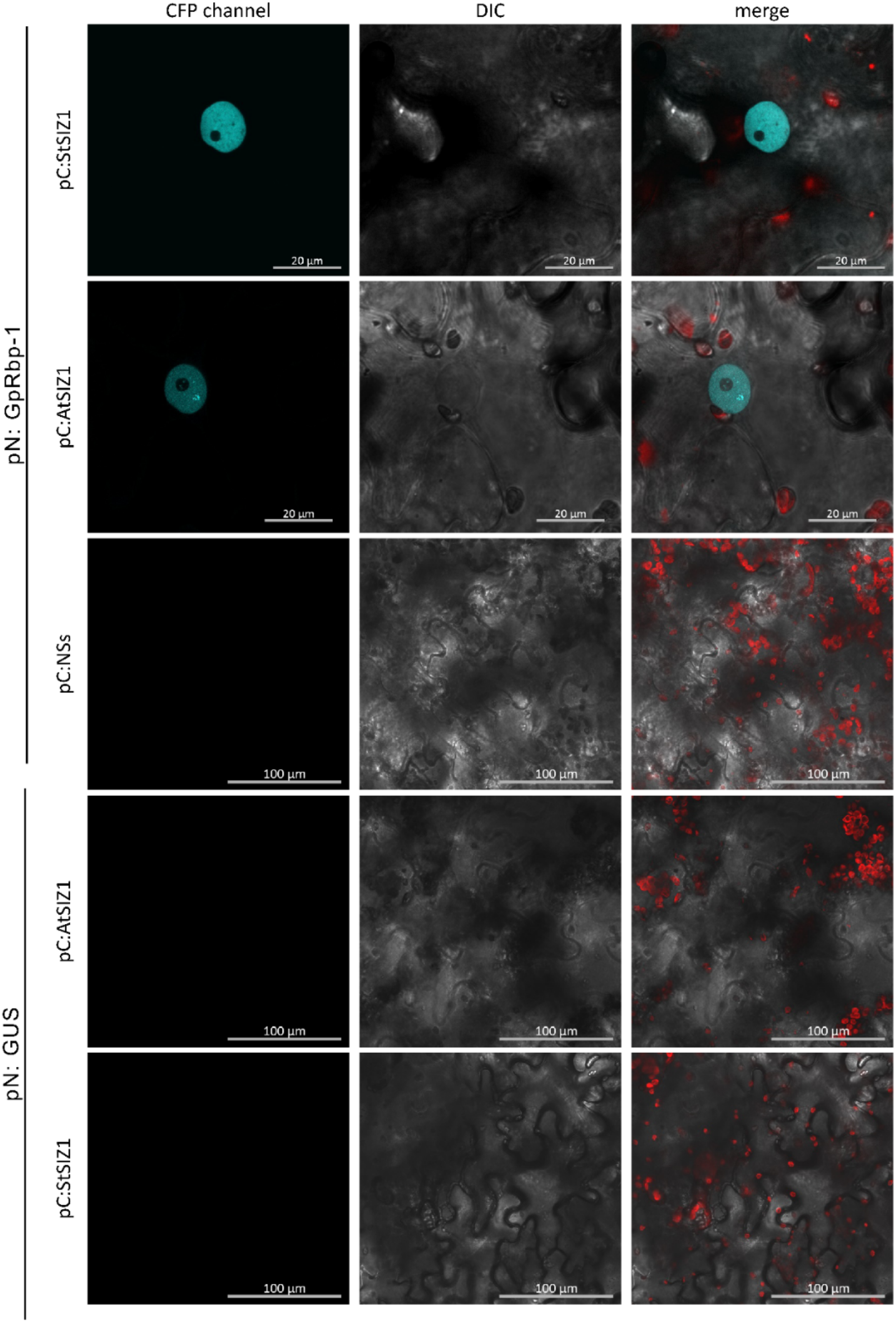
GpRbp-1 interacts with StSIZ and AtSIZ1 in the nucleus *in planta.* Bimolecular fluorescence complementation of N and C-terminal regions of SCFP3a. SCFP3A amino acids 1-173 were fused to GpRpb-1 (pN:Rbp1) and SCFP3A amino acids 156-239 were fused to StSIZ1(pC:StSIZ1) or AtSIZ1(pC:AtSIZ1) were coinfiltrated to *N. benthamiana* leaves. Co-expression of pN:EV or pC:EV were used as negative controls. The CFP emission channel is shown in blue, light emission in white in the differential interference contrast (DIC) channel and chloroplast auto-fluorescence is shown in red in the merge channel. Results are representative of 2 biological repeats and all co-infiltrations contain the silencing suppressor p19.

Having confirmed that GpRbp-1 targets StSIZ1 in the nucleus, we wondered if GpRbp-1 was also able to interact with distant homologues of SIZ1 from plant species which were also infected by cyst nematodes like the model species Arabidopsis. To investigate if GpRbp-1 was able to interact with AtSIZ1, we performed similar BiFC assays. N-terminally tagged AtSIZ with the C-terminal half of SCF3A (pC:AtSIZ1) was transiently co-expressed with pN:Rbp-1 in leaves of *N. benthamiana.* The fluorescent signal characteristic of SCF3A was reconstituted when pN:Rbp-1 was co-expressed with pC:AtSIZ1, but not when co-expressed with the negative control pC:NSs or pN:GUS in case of pC:AtSIZ (**Fig. 4**). The re-constituted signal indicating the interaction of pN:Rbp-1 and pC:AtSIZ1 was only visible in the nucleus of co-transformed cells. This shows that effector GpRbp-1 is also able to interact with AtSIZ1 *in planta* and that this interaction was limited to the nuclear cavity.

### SUMO-E3 ligase SIZ1 is involved in cyst nematode infection of *A. thaliana*

Based on the selective interaction of GpRbp-1 with both StSIZ1 and AtSIZ1, the role of SIZ1 in nematode infection was further tested *A. thaliana* and the beet cyst nematode *Heterodera schachtii* as a model system. First, we investigated the expression of *AtSIZ1* during cyst nematode infection. We measured the expression of *AtSIZ1* by quantitative RT-PCR in whole-roots of *H. schachtii* or mock-inoculated plants (Columbia-0) at 2, 4, 10, and 14 days postinoculation (dpi). No differential expression of the *AtSIZ1* transcript upon nematode infection was detected (**Suppl. Fig. 4**). From these data, we concluded that the *AtSIZ1* gene expression is not differentially regulated during cyst nematode infection which is consistent with a regulatory role in post-translational modification of proteins required for nematode parasitism.

To investigate if SIZ1 was involved in nematode infection we challenged *in* vitro-grown *siz1-2* knockout *A. thaliana* mutant with *H. schachtii.* These mutant lines carry an independent homozygous T-DNA insertion at different sites of exon 16 of the *AtSIZ1* gene resulting in a knock-out of SIZ1 (Lee *et al*., 2006; Miura *et al*., 2005). The homozygosity of the T-DNA insertion was verified using PCR primers designed with the iSect tool from the SALK Institute Genomics Analysis Laboratory (**Suppl. Table 6**). To examine the importance of AtSIZ1, the total number of nematodes infecting the roots of the mutant and wild-type control (Col-0) were counted at 14 dpi. Furthermore, we discriminated between adult female and male nematodes, as this indicates the nutritional quality of the established infection sites (Anjam *et al*., 2018; Trudgill, 1967). We observed a significant decrease of 36% (one-way ANOVA, p<0.001) in the total number of nematodes infecting the roots of *siz1-2* mutant as compared to the wild-type plants, respectively (**Fig. 5; Suppl. Fig. 5**). In addition, the number of female nematodes present in the roots of *siz1-2* was reduced by 49% in the mutant as compared to wild-type plants (one-way ANOVA, p<0.001). A similar effect was observed for the number of males, where a decrease of 29% in *siz1-2* plants as compared to the wild-type was found (one-way ANOVA, p<0.001) (**Suppl. Fig. 5**). Under *in vitro* growth conditions we did not observe an aberrant growth phenotype of the roots in the *siz1-2* seedlings, which was consistent with previous reports (Castro *et al*., 2015; Catala *et al*., 2007; Miura *et al*., 2011). Hence, the reduction in susceptibility could be attributed to the *siz1-2* mutation and not to differences in the mutant root systems. Therefore, we concluded that SIZ1 plays a role in the susceptibility of *Arabidopsis* to infection by cyst nematodes.

**Figure 5.**
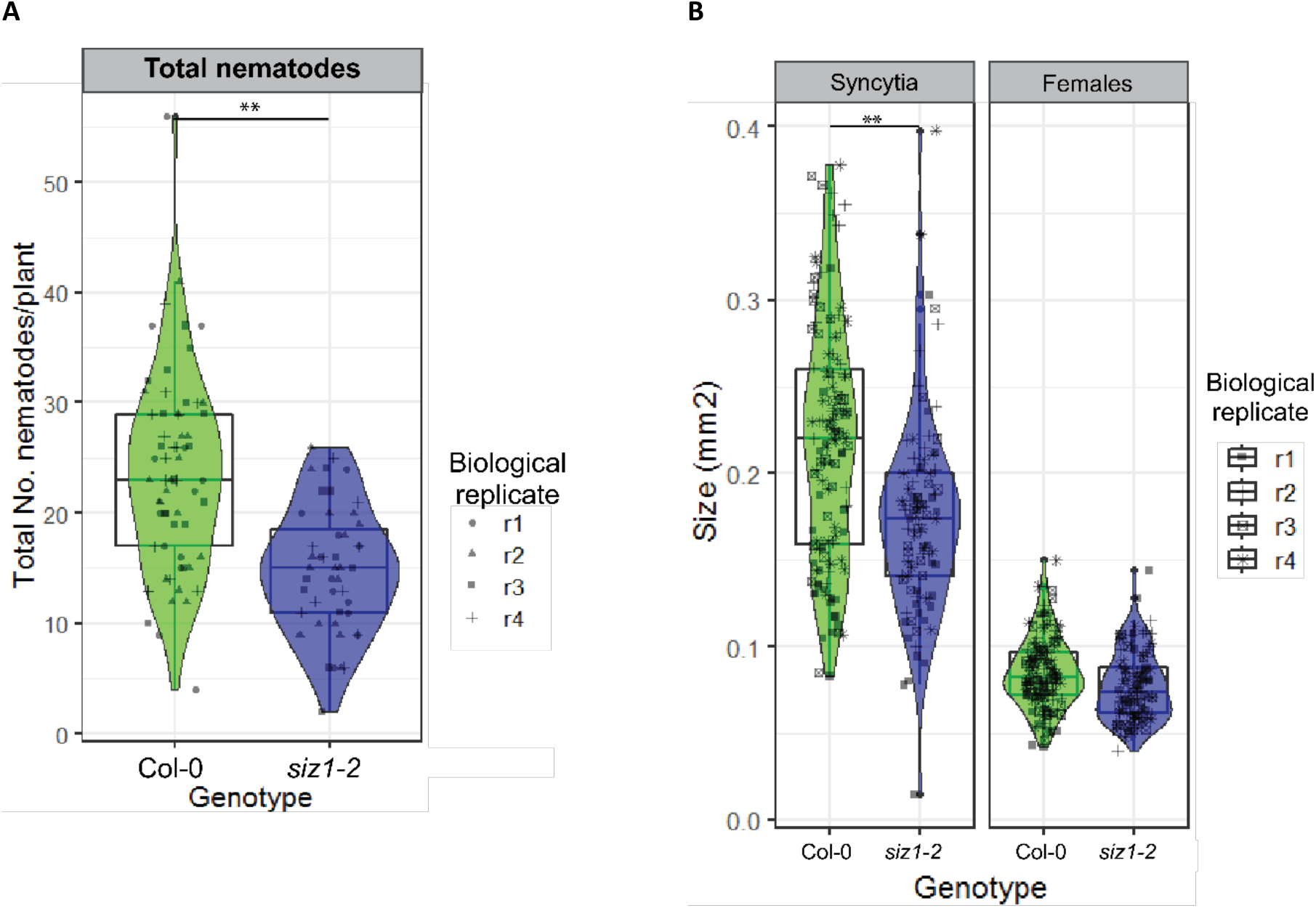
Knock-down of SIZ1 reduces nematode infection in *A. thaliana.* **A)** Average number of nematodes per plant in the roots of *Arabidopsis thaliana SIZ1-2* (n=47) and the background and wild type Columbia 0 (Col-0; n= 65) at 14 dpi. **B)** Average surface area (mm^2^) of female nematodes and syncytia present in the roots of *siz1-2* (n=94) and Col-0 (n=128) after 2 weeks of infection. Whiskers indicate the quartile (25 or 75%)+/− 1.5x interquartile range and violin plots describe the distribution of all data points. Stars indicate a significant statistical difference as determined by a one-way ANOVA (**p<0.001). Results are combined measurements from 4 independent biological repeats, using a fixed effects model.

Additional evidence was obtained by investigating the effect of SIZ1 on the size of the feeding site and growth of female nematodes infecting the roots of Arabidopsis. To this end, we measured the surface area of syncytia and females visible in the roots of *siz1-2* and wildtype Arabidopsis after two weeks of infection. The size of the syncytia induced by *H. schachtii* in *siz1-2* was 20% smaller as compared to the wild-type (one-way ANOVA, p<0.001) (**Fig. 5**). The size of the females established in *siz1-2* plants was not significantly different as compared to those developing on wild-type Arabidopsis plants (one-way ANOVA, p=0.335) (**Fig. 5**). Together, these results suggest that SIZ1 may not only contribute to the control of the overall infection rate, but might also influence the expansion of the permanent feeding sites of cyst nematodes in Arabidopsis.

### AtSIZ1 may contribute to plant defence to cyst nematodes

To further understand the role of SIZ1 during nematode interactions we performed a whole transcriptome analysis of *siz1-2* and wild-type Arabidopsis roots infected with beet cyst nematodes. To uncouple the effects of a mutated genotype and infection, we isolated whole root RNA from mock-inoculated and cyst nematode inoculated plants for both *siz1-2* and the wild-type plants, 7 days after inoculation (n = 3 replicates for each sample). We observed the overall expression of 13,114 genes in all 12 samples, and principal component analysis (PCA) showed a clear distinction between the *siz1-2* mutant and wild-type (first principal component, 27.1% of variance), and non-infected versus *H. schachtii* infected (second principal component, 13.4% of variance) (**Fig. 6**) as expected. Interestingly, the non-infected *siz1-2* and infected *siz1-2* samples cluster closer together on the 2^nd^ principal component axis than the non-infected and infected wild-type. This clustering indicates that *siz1-2* plants show less difference upon infection, the impact of the cyst nematodes on the transcriptome is smaller in the *SIZ1* mutant than in the wild-type (**Fig. 6**). The infection-like transcriptional profile of *siz1-2* is likely reflecting the auto-immune phenotype previously reported for *siz1-2* and the role of *SIZ1* as negative regulator of SA-mediated stress responses (Lee *et al*., 2006).

**Figure 6.**
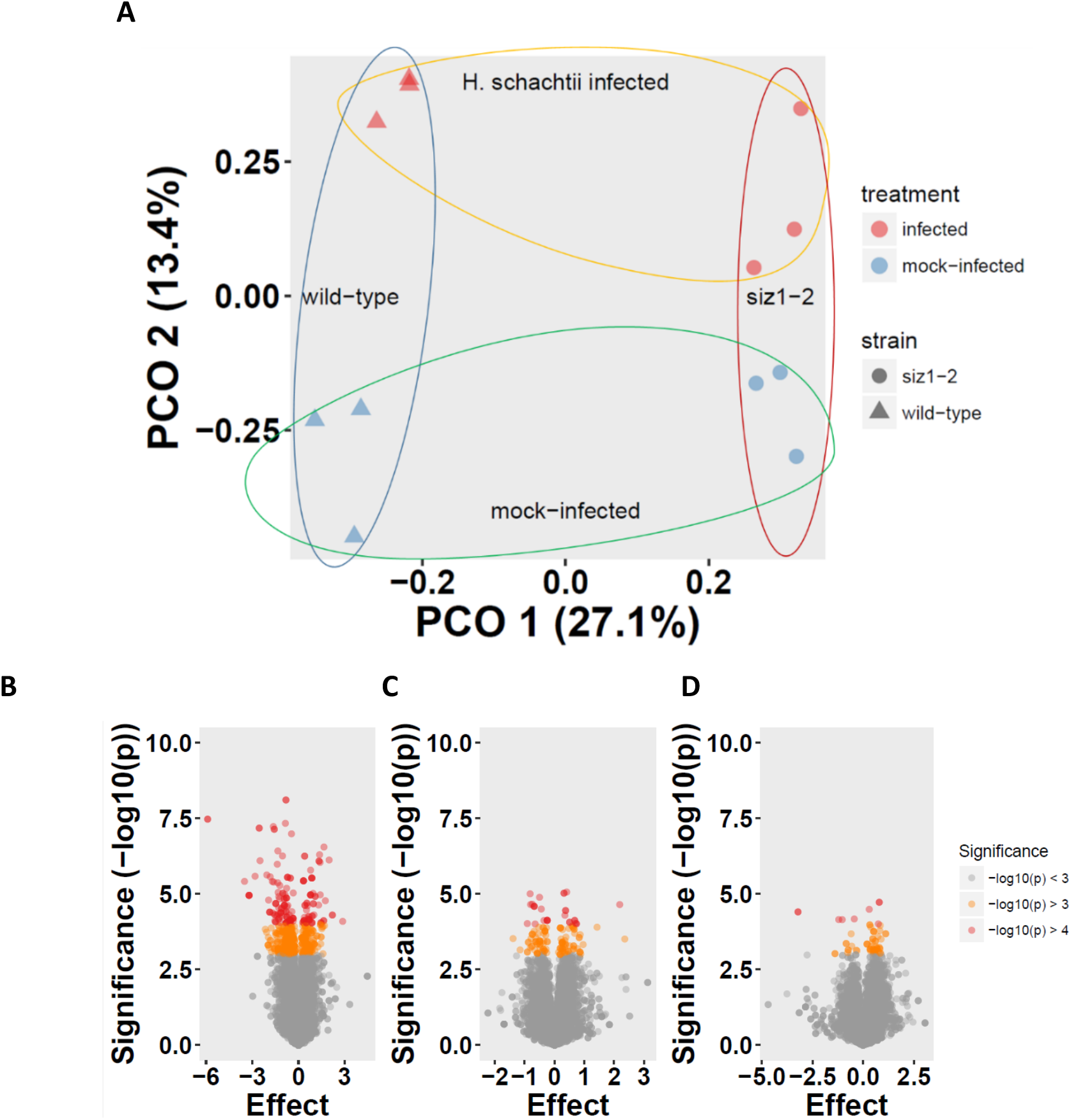
The mutation in *siz1-2* plants has a stronger effect in gene expression than nematode infection. **A)** Principal component analysis of gene expression profiles of *siz1-2* and wild-type Arabidopsis 7 days after (mock) infection with cyst nematodes. The first principal component PCO 1 captures 27.1% of the variation and separates the Arabidopsis seedlings by genotype. The second principal components PCO 2 captures 13% of variation and separates infected from uninfected samples. **B-D)** Volcano plots of differential gene expression as determined by total RNA RNAseq. The x-axis (effect) shows the relative expression of genes. The values on the y-axis reflect the significance of the differences in expression levels. The colours provide a visual aid for the thresholds mentioned in the legend. **B)** Genes differentially regulated in roots of *siz1-2* and wild-type Arabidopsis, irrespective of infection. **C)** Genes differentially regulated by nematode infection, irrespective of genotype. **D)** Genes differentially regulated by infection with *H. schachtii* in roots of *siz1-2* vs wild-type *Arabidopsis* plants, 7 days after inoculation.

To uncover which genes contributed to the separation of the samples in the PCA, we used a linear model to find 171 genes contributing to the difference between *siz1-2* and wildtype, 29 genes between non-infected and *H. schachtii* infected plants, and 9 genes between *siz1-2* and wild-type upon infection (interaction) (linear model, p < 0.0001; FDR_genotype_ = 0.011, FDR_infection_ = 0.064, FDR_interaction_ = 0.131; **Fig. 6; Suppl. Table 3; Suppl. Table 4**). To examine the hypothesis that nematodes manipulate a specific molecular pathway through SIZ1, we evaluated the gene ontology (GO) annotations of the 9 genes that are differentially regulated by the combination of mutant genotype and infection (**Suppl. Table 5**). Genes involved in control of the cell cycle (e.g. CYC B2;2), defence (e.g. BAP2) and protein transport (e.g. SLY1) were found in this differentially regulated group. Nevertheless, the small number of genes affected by the combination of the *siz1-2* mutation and the nematode infection (interaction) limits further interpretation of the molecular processes or pathways that nematodes may manipulate through SIZ1. In addition, the strong transcriptional differences between wild-type and *siz1-2* plants prior to infection, supported the notion that a mutation in SIZ1 induces drastic changes in the plant, even more so than infection with cyst nematodes.

To understand further the specific genes that were affected by the *siz1-2* mutation, we performed a gene-enrichment analysis. Thirty gene ontology (GO) terms are significantly upregulated in the *siz1-2* mutant, including “located in cell wall”, “involved in abiotic or biotic stimulus”, “cellular response to ethylene stimulus”, and “functions in carbohydrate binding” (**Suppl. Table 5**). In contrast, seven gene ontology categories are significantly upregulated in the wild-type, including “functions in sequence-specific DNA binding”, “located in extracellular region”, “involved in response to cold”, and “functions in sequence-specific DNA binding” (**Suppl. Table 5**). It is worth noting that the GO term most significantly upregulated in *siz1-2* plants is “involved in cellular response to ethylene stimulus”. Likewise, the GO term most significantly upregulated in wild-type plants is “functions in sequence-specific DNA binding” (**Suppl. Table 5**). These findings further support the existence of an “infection-like” transcriptional state of the *siz1-2* plants as detected with the PCA analysis due to constitutive activation of defence pathways as previously reported.

## Discussion

GpRbp-1 is an effector secreted by the potato cyst nematode *G. pallida* during the onset of parasitism, presumably to promote nematode virulence. To characterise the virulence role of this effector we aimed to identify the host proteins targeted by GpRbp-1. We found that GpRbp-1 interacts specifically in yeast and *in planta* with the potato homologue of the SUMO E3 ligase SIZ1. Furthermore, evidence from live cell imaging indicates that this interaction was limited to the nucleus of the cell, where GpRbp-1 co-localizes with StSIZ1. In addition, we evaluated the role of SIZ1 during nematode parasitism in the Arabidopsis - *H. schachtii* model system. *In vitro* infection studies in the Arabidopsis *siz1-2* mutant show that SIZ1 contributes to infection by cyst nematodes presumably as a negative regulator of plant defence. These results suggest that GpRbp-1 may target SIZ1 to repress plant immunity. This provides therefore evidence for the targeting of the master regulator SIZ1 by a pathogen effector to promote its virulence.

Likewise, the involvement of SUMOylation in plant-nematode interactions had not been described previously to the best of our knowledge. In contrast, SUMOylation has been shown to play a role in virulence of other plant pathogens. For instance, proteins from the SUMO machinery are transcriptionally regulated during infection by *Phytophthora infestans* in potato (Colignon *et al*., 2017). Also, interaction of replication protein AL1 from the geminivirus Tomato Golden Mosaic Virus and the SUMO E2-conjugating enzyme SCE1, is required for viral infection in *N. benthamiana* (Castillo *et al*., 2004; Sanchez-Duran *et al*., 2011). Furthermore, the effector XopD from *Xanthomonas euvesicatoria* has SUMO-protease activity, and catalyses the removal of SUMO from the tomato transcription factor SlERF4 to supress ethylene-mediated immune responses (Kim *et al*., 2013).

The reduced susceptibility of *A. thaliana* mutants *siz1-2* to cyst nematode infection (**Fig. 5**) likely reflects the role of SIZ1 as a negative regulator of plant immunity (Gou *et al*., 2017; Hammoudi *et al*., 2018; Lee *et al*., 2006; Niu *et al*., 2019). The *Arabidopsis SIZ1-2* mutant is characterised by a dwarf phenotype associated to increased levels of salicylic acid (SA) (Lee *et al*., 2006). This increased SA production in *siz1-2* is also associated to an upregulation of pathogenesis-related (PR) genes such as *PR1* and *PR5* (Lee *et al*., 2006). Additionally, the mutation in *siz1-2* confers resistance to the bacterial pathogen *Pseudomonas syringae* p.v *tomato (Pst*) (Lee *et al*., 2006). In contrast, the susceptibility of *siz1-2* to the fungal pathogen *Botrytis cinerea* is comparable to that of the wild-type (Lee *et al*., 2006). Therefore, Lee and co-workers (Lee *et al*., 2006) proposed that SIZ1 regulates immunity mediated by SA to biotrophic pathogens like *P. syringae,* independent of the jasmonic acid (JA) signalling pathway induced by the necrotrophic pathogen *B. cinerea.* Interestingly, cyst nematodes are also biotrophic pathogens, which trigger local and systemic SA-mediated plant defence responses upon root invasion (Kammerhofer *et al*., 2015; Lin *et al*., 2013; Nguyen *et al*., 2016; Wubben *et al*., 2008; Youssef *et al*., 2013). Together, these results suggest that SIZ1 may be required for basal resistance to biotrophic pathogens with different modes of infection above- and belowground.

In line with the auto-immune phenotype of the *siz1-2* mutant (Lee *et al*., 2006), our gene enrichment analysis shows that stress-related genes are differentially regulated in the *siz1-2* mutant in the absence of nematode infection (**Fig. 6; Suppl. Table 4**). It should be noted that we did not find elevated levels of *PR-l* or *PR-5* SA-responsive genes in the roots of *siz1-2* Arabidopsis. This observation may be due to the differences in growth conditions that may repress accumulation of SA or specific events in the SA-responsive pathway, or to different expression patterns of *PR* genes in roots and shoots. In addition, very few genes were differentially regulated in response to the infection of the mutant, suggesting that nematode infection has a relatively minor effect on transcriptional regulation of plant roots, as compared to the mutation alone (**Fig 6; Suppl. Table 4**). Therefore, our RNA-Seq data supports the hypothesis that the auto-immune phenotype of the *siz1-2* mutant underlies the mutant’s reduced susceptibility to infection by cyst nematodes. From this, a model can be inferred in which the immune-repressive function of SIZ1 in SA-mediated defence responses is enhanced by nematode effectors leading to an increase in the susceptibility of host plants to cyst nematode infections.

In plants, a zinc-finger motif within the SP/RING domain is largely responsible for the nuclear localization of AtSIZ1 as well as the regulatory role of SA accumulation and SA-dependent phenotypes (e.g. dwarfism, resistance to pathogens and thermotolerance) (Cheong *et al*., 2009). Interestingly, GpRbp1 interacts with a protein fragment comprising this domain of SIZ1, suggesting that it may affect the SUMO E3 ligase activity of SIZ1. Furthermore, the nuclear co-localization and interaction of GpRbp-1 and StSIZ1 seems to follow specific substructures within the nucleus (**Figs. 3 and 4**). This may point at targeting of StSIZ1 by GpRbp-1 to modify or modulate the nuclear activity of StSIZ. In yeast, the RING domain is necessary to recruit the E2-SUMO complex into a complex with its substrate (Yunus and Lima, 2009). Hence, targeting of the SP-RING finger domain of SIZ1 by GpRbp-1 most likely compromises these SA-related phenotypes. In turn, the characteristic hypervariable region of GpRbp-1 may function as a binding platform to facilitate the targeting of SIZ1 (Diaz-Granados *et al*., 2016; Rehman *et al*., 2009).

Additionally, the gene ontology “involved in cellular response to ethylene stimulus” was the enriched category with the highest statistical support. From the differentially regulated genes, LEC1 (lectin-like protein; AT3G15356), FRD3 (Ferric reductase defective 3; AT3G08040), and RBK1 (ROP binding protein kinase; AT5G10520) are grouped in the “response to ethylene stimulus” ontology (**Suppl. Table 4**). LEC1 has been shown to be transcriptionally regulated in response to several stimuli, including the fungal elicitor chitin and mechanical wounding. The response of LEC1 to chitin is also found in ethylene/jasmonate (ET/JA)-insensitive mutants, suggesting that LEC1 is involved in ET/JA-dependent and independent cellular responses (Seoung Hyun *et al*., 2009). This finding could indicate that, opposite to previous hypotheses (Lee, 2006), SIZ1 may be involved in regulation of the JA defence pathway through modulation of the JA/ET branch (Pieterse *et al*., 2012). Moreover, in plant-nematode interactions ethylene can act as a modulator of SA-immunity or as a regulator or cytokinin-dependent susceptibility, and these roles are determined by the activation of specific ethylene receptors (Piya *et al*., 2019). Therefore, it remains to be determined if SIZ1 acts solely as a regulator of SA-mediated plant immunity in plant nematode interactions.

SIZ1 may also act as regulator or hormone-dependent metabolic processes that influence susceptibility to nematodes. The decrease in the size of the specialised feeding sites (syncytia) induced by nematodes in *siz1-2* plants points to a role of SIZ1 in expansion of nematode-induced feeding sites (**Fig. 5**). Additionally, the genes differentially regulated by the *siz1-2* mutation in combination with cyst nematode infection may have further roles as regulators of feeding site formation. For example, in addition to ethylene responsiveness, FRD3 is also involved in nutrient homeostasis and iron uptake (Xing *et al*., 2015). And in turn, RBK1 has been shown to be regulated by pathogen infection (Molendijk *et al*., 2008), but also to be implicated in auxin-mediated cell expansion (Enders *et al*., 2017). Both nutrient uptake and cell expansion processes are relevant in the context of nematode feeding sites (reviewed in (Kyndt *et al*., 2013)). Finally, SIZ1 represses the characteristic root morphological adaptations to phosphate starvation, through the control of auxin patterning (Miura *et al*., 2011; Miura *et al*., 2005). Auxin transport and signalling are involved in the proper formation of syncytia, presumably by its role as regulator of plant organogenesis (reviewed in (Gheysen & Mitchum, 2019; Ng *et al*., 2015)). These findings illustrate how the regulatory network of SIZ1 becomes intricate, with effects on different plant hormones that are implicated in immune responses as well as in cellular modifications related to nematode infection. Ultimately, elucidating the mechanism of action of SIZ1 would require the identification of the molecular targets that are regulated by SIZ1 in plant-nematode interactions. Nevertheless, around 600 proteins are predicted as potential SIZ1-dependent SUMO targets in plants (Rytz *et al*., 2018). Consequently, a definite mode of action of SIZ1 in plant-nematode interactions will require further dissection through a combination of genetic, biochemical and *in vivo* assays.

Preliminary data also provide evidence for a complementary hypothesis where GpRbp-1 may modulate host cellular processes by recruiting the SUMO machinery of the host. Here, we could show that GpRbp-1 interacts in BiFC with other components of the SUMO machinery, namely SUMO1, 3 and 5 (SUM1, SUM3, SUM5) and the E2 conjugating enzyme SCE1 (**Supp. Fig. 6**). Interestingly, GpRbp-1 interacts with the SUMOs and SCE1 in the nucleus and cytoplasm, where these proteins localize (**Suppl. Fig. 6**) (Mazur *et al*., 2019; Xiong & Wang, 2013). In yeast and mammals, multi-protein complexes including SUMO, E2 (Ubc9), and E3s (PIAS/Nup358) (Mascle *et al*., 2013; Reverter & Lima, 2005) are required for SUMOylation and the ensuing transcriptional regulation activities of UBC9 and SUMO1 (Mascle *et al*., 2013; Reverter & Lima, 2005). Similarly, in *Arabidopsis* SUMO, SCE1 and SIZ1 form a ternary complex that is recruited to nuclear bodies (NBs) where COP1 is SUMOylated to regulate the response of the plant to darkness and temperature (i.e. skoto- and thermo-morphogenesis) (Kim *et al*., 2016; Lin *et al*., 2016; Mazur *et al*., 2019; Osterlund *et al*., 2000; Seo *et al*., 2003; Yang *et al*., 2005). Conceivably, the virulence role of GpRbp-1 may be exerted through an influence on the SUMO-SCE1-SIZ1 tertiary complex, for example by stabilisation. In support of this notion, our co-immunoprecipitation assays suggest that intermediate compounds present in a larger complex with StSIZ1frag14 and StSIZ1frag83 are co-pulled down specifically by GpRbp-1 (**Fig. 2**). The nature of the complex co-pulled down along with StSIZfrag14 and StSIZ1frag83 by GpRbp-1 the remains to be established.

An alternative explanation is that GpRbp-1-like effectors may recruit the SUMO complex to achieve SUMOylation inside the host cells for full functionality as an effector. Conceivably the SUMOylation of GpRbp-1-like effectors may enhance their stability by competing with ubiquitination (Zheng *et al*., 2018) or modify their binding patterns (Guo & Sun, 2017; Hansen *et al*., 2017). This hypothesis is supported by the prediction of consensus SUMO-acceptor ΨKxE) and SUMO-interaction motifs (SIM) in GpRbp-1 (**Suppl. Fig. 7**), indicating that host-mediated SUMOylation may be relevant for its functioning as an effector. Two SIM are predicted in the N-terminal half of GpRbp-1, in one region that is unique to GpRbp-1 and another present in several Rbp and SPRYSEC sequences (Diaz-Granados *et al*., 2016). The first SIM, unique to GpRbp-1 falls within a region with low confidence for modelling, so it is difficult to predict in what region of the GpRbp-1 structure it is located. The second SIM (SIM2) localizes to a β-sheet present in the core of the β-sandwich structure that is predicted for GpRbp-1 (**Suppl. Fig. 7**). In addition, two ΨKxE SUMOylation sites are predicted, one inverted in the conserved core of the SPRY domain (ΨKxE_nverted) and one in the C-terminus in a motif present only in GpRbp-1 and Rbp-1 from *G. mexicana* (ΨKxE_2) (Diaz-Granados *et al*., 2016) (**Suppl. Fig. 7**). The “conserved” SIM2 and (ΨKxE inverted) sites reside in the β-sheet core of GpRbp-1, and most likely form a binding pocket for SUMO, whereas the unique ΨKxE_2 site resides in a C-terminal α-loop and is likely exposed to the solvent (**Suppl. Fig. 7**). None of these motifs contain residues reported to be under positive selection in *G. pallida* field populations (Carpentier *et al*., 2012). In this scenario, the lack of SIZ1 in *SIZ1-2 Arabidopsis* impedes an efficient functioning of GpRbp-1 homologues from *H. schachtii* as a virulence factor. Although the functional homolog of GpRbp-1 of *H. schachtii* is not known, we assume that similar proteins may exist based on the existence of GpRbp-1-like gene transcripts (Fosu-Nyarko *et al*., 2016) which may exert a similar function in plant parasitism. Moreover, different cyst nematode species share the same mode of parasitism, which results in the formation of typical feeding structures. Therefore, we expect that the underlying molecular mechanisms are conserved among host plant species. This is supported by our observation that GpRbp-1 can also interact with AtSIZ1 (**Fig. 4**), suggesting that SIZ1 may be a conserved target of cyst nematodes.

## Experimental procedures

### Yeast two-hybrid – library screen

The prey *G. pallida-* infected potato library was generated by Dual Systems Biotech (Switzerland) from grinded roots of potato *(Solanum tuberosum* ssp. *andigena*) genotype SH infected with juveniles of *G. pallida* population Pa3-Rookmaker. Potato plantlets were grown on 16 cm square plates containing B5 medium at 20°C in 16 h light/ 8 hours dark conditions. Two weeks after transplant, plantlets were inoculated with ~200 juveniles. Infected roots were collected at 2, 3, 7, 9, 12 and 14 dpi and grinded in liquid nitrogen before shipping. Poly (A) tailing and total RNA isolation were performed by Dual Systems Biotech (Switzerland) and a cDNA library consisting of 3,85×10^6^ clones with an average insert size of 1.13Kb was constructed. The library yeast two-hybrid screen was performed by Dual Systems Biotech (Switzerland) using the DUALhybrid vector system.

### Cloning

For co-immunoprecipitation, the interacting fragments StSIZ1fragDS14 and StSIZ1fragDS83 were excised from the pGAD-HA prey vector using NcoI and XhoI restriction sites. The NcoI-XhoI fragment includes the StSIZ1fragDS14 and StSIZ1fragDS83 with an N-terminal hemagglutinin (HA) tag. The NcoI-XhoI fragment was inserted into vector pRAP digested with NcoI and Sail. In the vector and additional four units of N-terminal HA tag are fused to the HA-StSIZ1fragDS14 and HA-StSIZ1fragDS83. The fusion cassette was digested with PacI and AscI and ligated into vector pBIN. In pBIN, the HA-StSIZ1fragDS14 and HA-StSIZ1fragDS83 fusion is under the control of the 35S constitutive promotor. GpRpb-1 version 1 from virulent population Rookmaker (Rookl) was tagged with an N-terminal fusion of 4 units on the c-Myc tag followed by a GFP (Myc4-GFP-GpRbp-1) and transferred by restriction enzyme cloning to the pBINPLUS binary vector (van Engelen *et al*., 1995). For microscopy studies, the full-length gene of StSIZ1 was obtained by synthetic gene synthesis (GeneArt) (Thermo Fisher Scientific, Waltham, Massachusetts) and cloned to appropriate pGWB vectors (Nakagawa *et al*., 2007) by Gateway cloning. The mCherry GpRbp-1 construct was generated by restriction cloning in pBINPLUS (van Engelen et al., 1995). BiFC constructs were generated by Gateway cloning to the pDEST-SCYCE(R)^GW^ and pDEST-SCYNE(R)^GW^ vectors (Gehl *et al*., 2009).

The full-length gene of StSIZ1 was obtained by synthetic gene synthesis (GeneArt) using the potato CDS transcript variant X3 (GenBank accession XM_006340080.2) into Gateway-compatible pMA vector. StSIZ1 was synthesized with an additional N-terminal BamHI site and a C-terminal PstI site in order to enable restriction cloning. The internal BamHI and PstI restriction sites were disrupted by introducing silent mutations. The codons were always replaced with ones with similar or higher usage frequency in *Nicotiana benthamiana* (Nakamura et al., 2000). Full-length SIZ1 N-terminally tagged with HA or GFP was obtained by gateway cloning to plant-expression vectors pGWB415 and pGWB425 respectively, (Nakagawa *et al*., 2007) for interaction and localization studies.

For bimolecular fluorescence complementation (BiFC), GpRbp1 was amplified by PCR and cloned into pDONR207 by a BP reaction, following the manufacturer’s instructions (Invitrogen). Expression clones of GpRbp1 were obtained by LR recombination into pDEST-SCYCE(R)^GW^ BiFC vector (Gehl *et al*., 2009). Similarly, expression clones of full-length SIZ1 (StSIZ1) were recombined by LR recombination (Invitrogen) into pDEST-SCYNE(R)^GW^ BiFC vector following the manufacturer’s instructions (Gehl *et al*., 2009). cDNA of SCE1, SUMO1, SUMO3 and SUMO5 from Arabidopsis was obtained from total RNA extracted with TRIzol reagent (Thermofisher), followed by first-strand cDNA synthesis using M-MLV reverse transcriptase (Promega) and random hexamers (Roche) according to the manufacturer’s protocols. Primer pairs containing attB1 and attB2 recombination sites (**Suppl. Table 6**), were used to amplify the coding sequences of SCE1, SUMO1, SUMO3 and SUMO5, respectively. The resulting PCR amplicons were recombined with the Gateway vector pDONR207 (Thermofisher) by BP reaction, following the manufacturer’s instructions. Subsequently, expression clones were generated by LR recombination into the pDEST-SCYNE(R)^GW^ BiFC vector following the manufacturer’s instructions (Gehl *et al*., 2009).

### Expression and detection of recombinant proteins

All proteins were co-expressed by Agrobaterium-mediated transient transformation of *Nicotiana benthamina* leaves. All co-expression assays are done together with the silencing supressor p19, with a final concentration of OD_600_=0.5. Total protein extracts were prepared by grinding leaf material in protein extraction buffer. For co-IP, pull-downs were performed using μMACS anti-c-MYC or anti-HA paramagnetic beads (Miltenyi, Bergisch Gladbach, Germany). Proteins were separated by SDS-PAGE on NuPage 12% Bis-Tris gels (Invitrogen, Carlsbad, California) and blotted to 0.45μm polyvinylidene difluoride membrane (Thermo Fisher Scientific). Immunodetection was performed with corresponding horseradish peroxidase-conjugated antibodies. Confocal microscopy was performed on *N. benthamiana* epidermal cells using a Zeiss LSM 510 confocal microscope (Carl-Zeiss, Oberkochen, Germany) with a 40X, 1.2 numerical aperture water-corrected objective.

### Confocal microscopy

*N. benthamiana* epidermal cells were examined using a Zeiss LSM 510 confocal microscope (Carl-Zeiss) with a 40X 1.2 numerical aperture water-corrected objective. For co-localization studies the argon laser was used to excite at 488 nm for GFP and chlorophyll, and the HeNe laser at 543nm to excite mCherry. GFP and chlorophyll emission were detected through a band-pass filter of 505 to 530nm and through a 650nm long-pass filter, respectively. mCherry emission was detected through a band-pass filter of 600 to 650nm. For BiFC the argon laser was used to excite at 458 nm for SCFP3A. SCFP3A emission was detected through a band-pass filter of 470nm to 500nm and chlorophyll emission was detected through a 615nm long-pass filter. We also used a CFP marker to calibrate the fluorescence excitation and emission for CFP.

### Plant material and nematode infection

Seeds of the homozygous *siz1-2* were kindly provided by Dr. H. van den Burg (Laboratory for Phytopathology, University of Amsterdam, the Netherlands). Col-0 N60000 wildtype seeds were obtained from the SALK homozygote T-DNA collection (Alonso et al., 2003).-For nematode infection, seeds were vapour sterilized and sown in modified KNOP medium (Sijmons *et al*., 1991) and grown at 25°C under a 16-h-light/8-h-dark cycle. 10 day-old seedlings were inoculated with 60-70 surface-sterilized *H. schachtii* infective juveniles. After 2 weeks of infection, the amount of nematodes present in the roots of Arabidopsis plants were counted visually and the size of females and syncytia were determined as described previously (Siddique *et al*., 2014). Statistical differences were estimated by one-way ANOVA (a=0,05), using the weighted-inverse variants to combine data from 4 biological replicates.

### RNA sequencing

#### Total RNA extraction

Seeds of *siz1-2* and Col-0 wild-type (N60000) were vapour sterilized and sown in modified KNOP medium (Sijmons *et al*., 1991) in 6-well cell culture plates (Greiner Bio-one). Seedlings were grown at 21°C under a 16-h/8-h light/dark regime. Two-week old seedlings were infected with approximately 180 surface-sterilized *H. schachtii* juveniles. One week after inoculation the complete root systems of ~18 *siz1-2* and Col-0 plantlets were harvested and snap-frozen. Root tissue was ground in liquid nitrogen and total RNA was extracted with the Maxwell^®^ 16 LEV plant RNA kit (Promega) in the Maxwell 16 AS2000 instrument (Promega), following the manufacturer’s instructions. Three biological replicates of ~18 plants/sample per condition were generated.

#### Count derivation and normalization

Quality checking, removal of adapter sequences, genome mapping and count derivation was performed by a custom-written pipeline. Briefly, read quality was assessed using FASTQC v0.11.5 (Andrews, 2014). Overrepresented adapter sequences, base pairs with a Q-value lower than fifteen in the 5’ or 3’ and reads shorter than 20bp were removed with cutadapt v1.16 (Martin, 2011). Reads were then mapped to the *A. thaliana* genome TAIR10 with Hisat v2.1.0v (Cheng *et al*., 2017; Kim *et al*., 2015);. In all samples, well above 85% of the reads mapped to Arabidopsis (**Suppl. Table 2**). Obtained SAM files were sorted and converted to BAM files with the help of samtools v1.6 (Li, 2011; Li *et al*., 2009). From these files FPKM counts of mapped sequences were derived by StringTie (Pertea *et al*., 2015).

The FPKM-transformed counts were imported in “R” (version 3.4.2, x64) and were log_2_ transformed by *C_i,j_* = log_2_(*FPKM_i,j_* + 1), where C is the log_2_-transformed FPKM value for gene i (one of 37217 unique transcripts) of sample j (one of three replicates of wild-type mock infected, wild-type infected, *siz1-2* mock infected, or *siz1-2* infected).

Subsequently, we selected only transcripts that were detected in all 12 samples (C > 0) for further analysis (representing 13114 unique genes). For principal component analysis, we also transformed to data to a log_2_-ratio with the mean, by 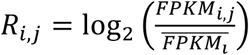 where R is the log_2_-ratio with the mean of transcript i of sample j, and 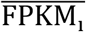 is the mean of the FPKM values for gene i.

Thereafter, both C and R values were batch-corrected by subtracting the mean difference of the batch from the total mean, as follows

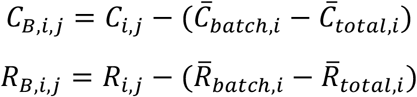

where C_B_ and R_B_ are the batch-corrected values of gene *i* of sample *j* and 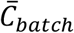 and 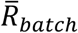 are the batch averages, and 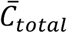 and 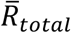 are the averages over the total.

#### Differential expression analysis

To understand the contributing factors underlying variance in the gene expression data, we first used a principal component analysis on R_B_ to understand the major sources of variance. Thereto we used the *prcomp* function in “R”.

We then applied a linear model to identify genes contributing to the genotype differences, the differences between mock-infection and infection, and the interaction between both variables, by solving *C_B,i,j_* = *G_j_* + *T_j_* + *G_j_*×*T_j_* + *e_j_* where C_B_ of gene *i* of sample *j* was explained over genotype (G; either wild-type or *siz1-2*) and treatment (T; either mock-infected or infected), the interaction between G and T, and an error-term (e). The significances were used to calculate a false discovery rate (FDR) using the *p.adjust* function in “R” (Benjamini & Hochberg, 1995). To make explanatory terms comparable, we applied a single significance threshold of p < 0.0001, which corresponded to a FDR of 0.011 for genotype, 0.064 for treatment, and 0.131 for the interaction between genotype and treatment.

The differentially expressed genes (p < 0.0001) were used in an enrichment analysis, as described before (Warmerdam *et al*., 2019). In short, enrichments were calculated by hypergeometric test, using the TAIR11 databases: Gene ontology, Gene ontology slim, gene classes, and phenotypes (Berardini *et al*., 2015; Lamesch *et al*., 2012), and the MapMan gene ontology database, based on TAIR10 (Thimm *et al*., 2004). We filtered groups were fewer than three genes overlapped, and selected significant enriched groups based on a correction for multiple testing (FDR).

#### Data availability

The data was submitted to ArrayExpress, under code E-MTAB-8193.

### Phylogenetic tree

Sequences were aligned in BioEdit v.7.2.6 (Hall, 1999) and a Bayesian tree was created using MrBayes v.3.2.26 (Ronquist *et al*., 2012). The data set was partitioned according to codon position and the analysis was run for 500,000 generations with a GTR + invariable sites + gamma substitution model using 4 MCMC chains and 4 parallel runs. After checking for conversion with Tracer v.1.7.1 (Rambaut *et al*., 2018), the burnin was set to 5,000 generations.

### GpRbp-1 modelling

The model of GpRbp-1 was built by remote homology modelling, using a similar workflow as previously described (Rehman *et al*., 2009). Briefly, an initial sequence analysis was performed by identifying specific sequence patterns signatures using InterProScan (Jones *et al*., 2014) and ScanProSite (De Castro *et al*., 2006). Consensus profiles for structural feature predictions were obtained using various methods, namely Jpred4 (Drozdetskiy *et al*., 2015), RaptorX-Property (Wang *et al*., 2016), SCRATCH (Cheng *et al*., 2005), PsiPred (Buchan & Jones, 2019) and Spider3 (Heffernan *et al*., 2017) (i.e. secondary structure, intrinsically disorder regions and relative solvent accessibility predictions).

The 3D model of GpRbp-1 sequence was built within the interval (aa 61-246) starting from the closest homologues with available crystal structures - namely the IUS-SPRY domain of human RanBP9 (Ran Binding Protein 9, PDB 5JI7) (Hong *et al*., 2016), mouse RanBP10 (Ran Binding Protein 10, PDB 5JIA) (Hong *et al*., 2016), and the SPRY domain of human SPRYD3 (SPRY Domain-Containing Protein 3, PDB 2YYO) (Kishishita *et al*., 2008), sharing 31.5%, 30.9% and 20.8% sequence identity respectively with GpRbp-1. The N-terminus region (aa 1 - 60), including the first extended PRY motif was not modeled, as no 3D templates with adequate homology were detected. The model was built using Modeller v9.20 (Webb & Sali, 2014) and further refined by iterative rounds local and global simulated annealing and energy minimization monitored with MolProbity (Williams *et al*., 2018) until convergence to a Molprobity quality score of 1.11Å from an optimal polypeptide path. The optimized model was further subjected to a 20 ns long molecular dynamics simulation for stability test. Molecular dynamics, simulated annealing and energy minimization stages were performed in explicit solvent in NAMD v2.12 (Phillips *et al*., 2005) using the CHARMM36M forcefield (Huang *et al*., 2017), TIP3 water molecules model and a fixed 0.15 M NaCl concentration, with the overall system size summing up to 27050 atoms (from which 2863 atoms correspond to GpRbp-1).

MD simulations were performed at 300 K constant temperature, using a 2 fs timestep, Particle Mesh Ewald full-system periodic electrostatics and periodic boundary conditions, Langevin temperature control and Nosé-Hoover Langevin piston for a constant 1 bar pressure control, as implemented in NAMD v2.12 (Phillips *et al*., 2005). The stability of the model was investigated by analyzing the potential energy, RMSD (root mean square deviations) and RMSF (root mean square fluctuations) along the simulation trajectory. All trajectory analyses and predictive model figures were obtained using VMD (Humphrey *et al*., 1996) and PyMOL v2.2.3 (DeLano, 2002).

## Acknowledgments

The authors would like to thank Casper van Schaik, Branimir Velinov and Jorrit Lind (Wageningen University) for technical assistance; Prof. Alvaro Muñoz (Johns Hopkins University) for advice on statistical analysis; Dr. Harrold van den Burgh (University of Amsterdam) for providing *siz1-2* seeds and AtSIZ1. E.C.M. and A.J.P. acknowledge support from UEFISCDI grant PN-III-ID-PCE-2016-0650. G.S., H.O., and A.D.G. acknowledge the Dutch Science Foundation NWO-ALW project 828.11.002 for funding and A. G., J.R and R.P. the Dutch Top Technology Institute Green Genetics for their support. This work benefited from interaction within the COST Action SUSTAIN FA1208.

## Supplementary Figures

**Supplemental Table 1.**
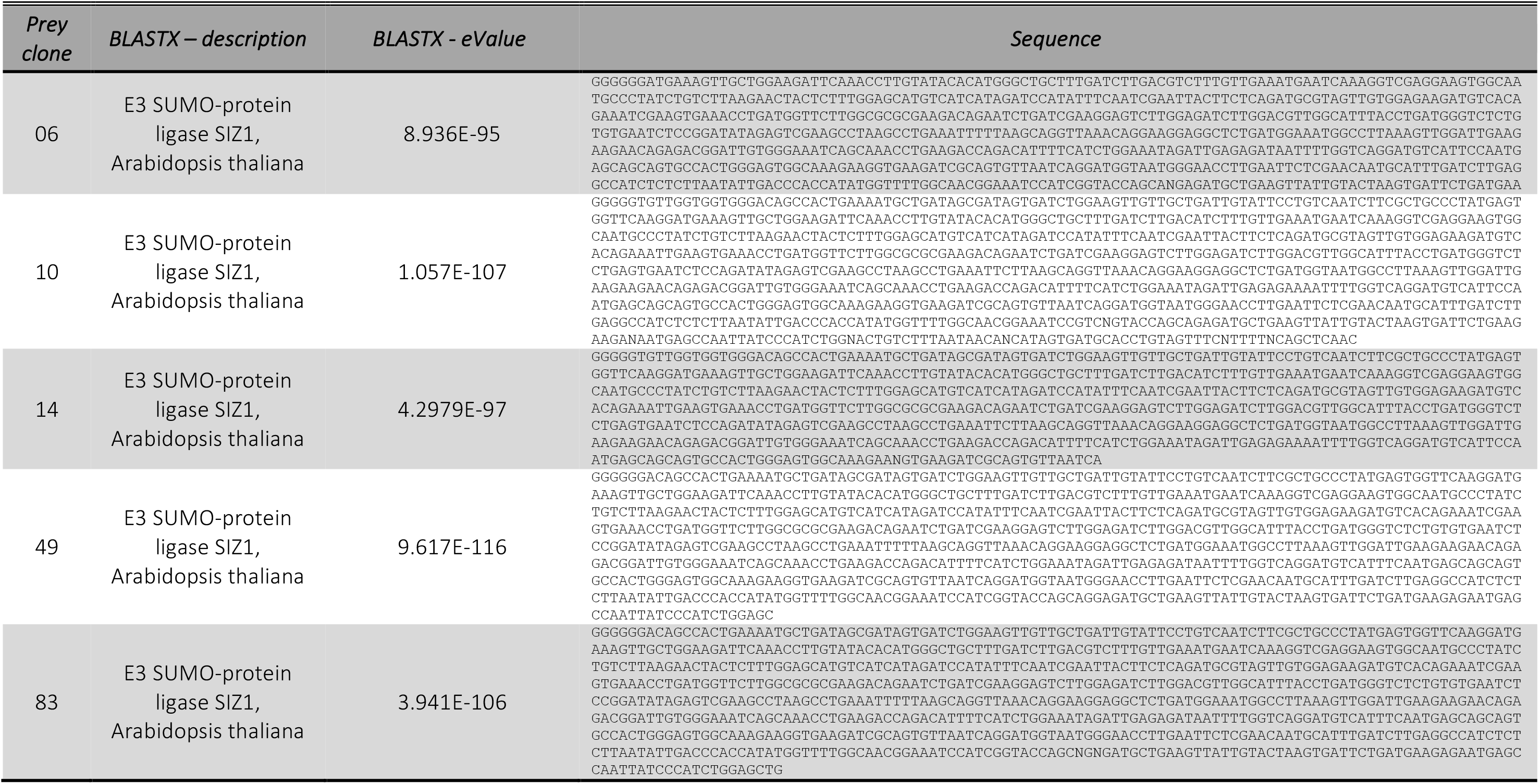
cDNA sequence of StSIZ1 fragments interacting with GpRbp-1 in yeast.

**Supplemental Figure 1.**
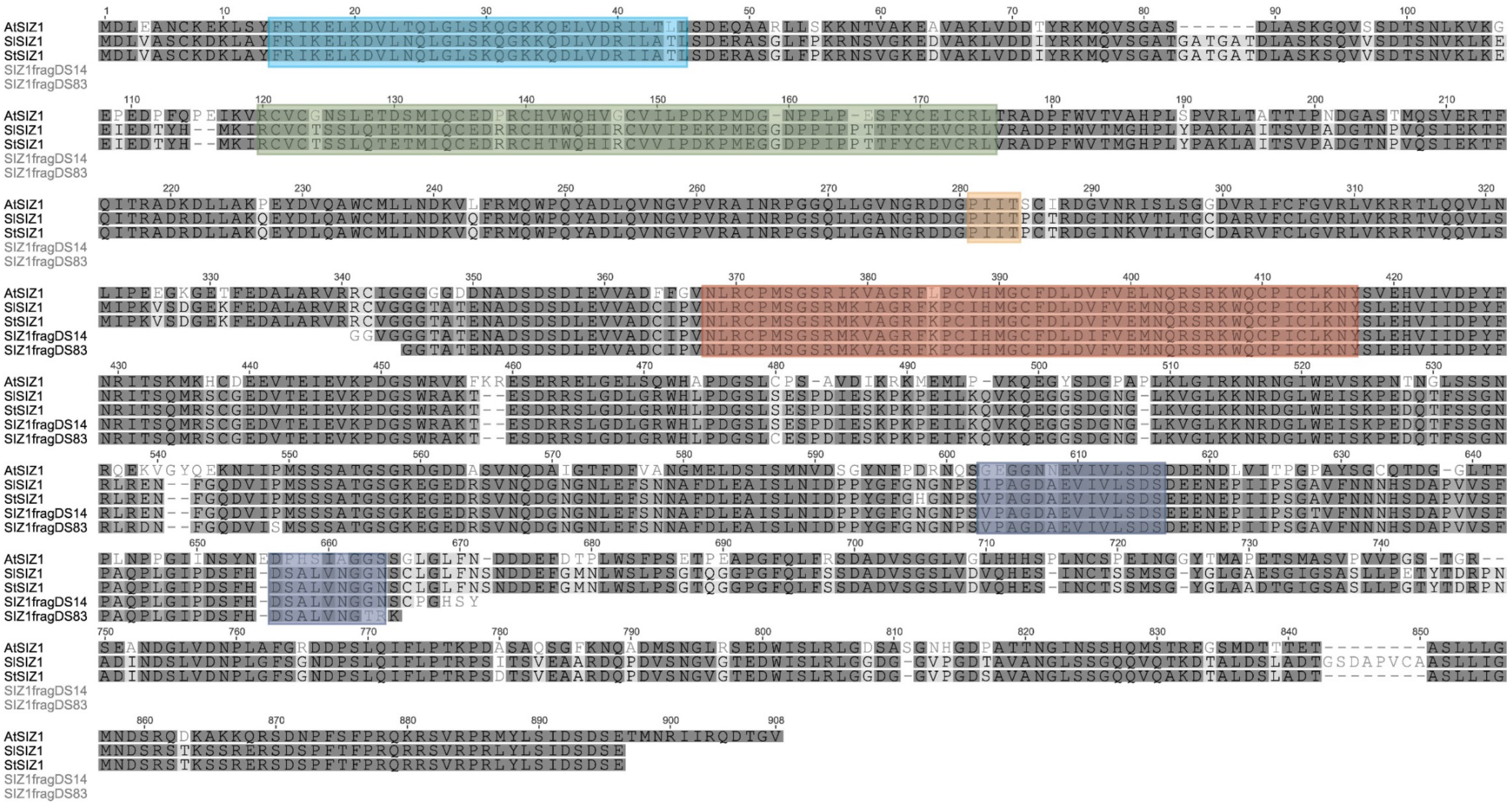
Full-length alignment of StSIZ1, AtSIZ1, SISIZ1, StSIZ1fragl4 and StSIZfrag83. Protein alignment made with the CLUSTALW plugin of Geneious (version 8.1.9) with cost matrix BLOSUM. The characteristic domains of SIZ1 are shown in coloured boxes: the SAP domain in light blue, the PHD domain in green, the PUT (PINIT) motif in orange, the SP-RING domain in red and the SIMs in dark blue. Full-length proteins were obtained from TAIR, AtSIZ (1009128193) or GenBank StSIZ1 (XP 006340142.1) and SISIZ1 (KP323389.1). The protein sequences encoded by the yeast-interacting fragments were obtained by automatic translation.

**Supplemental Figure 2.**
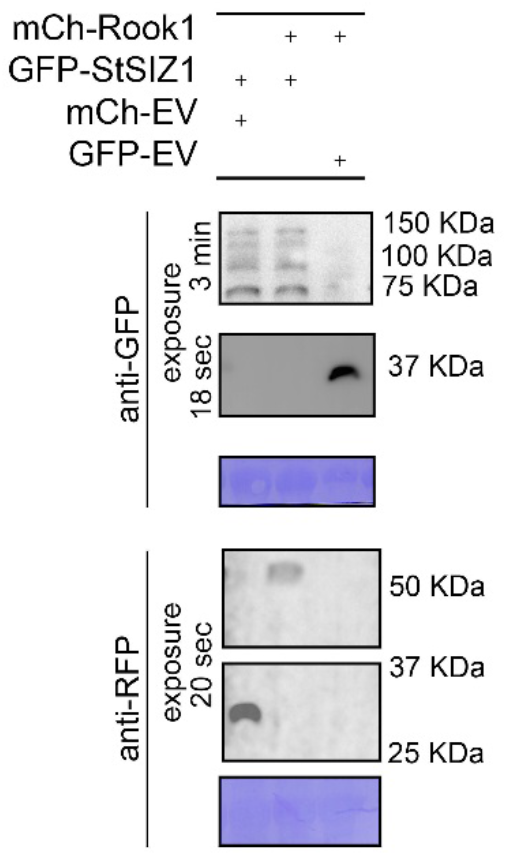
Fluorescent fusions of GpRbp81 and SIZ1 are expressed in leaves of *N. benthamiana.* Western blot detection of fusion of green fluorescent protein with StSIZ1 (GFP-SIZ1), red fluorescent protein mCherry (mCh-GpRbp-1) or GFP and mCh alone. Western blot is performed with anti-GFP and anti-RFP antibodies.

**Supplemental Figure 3.**
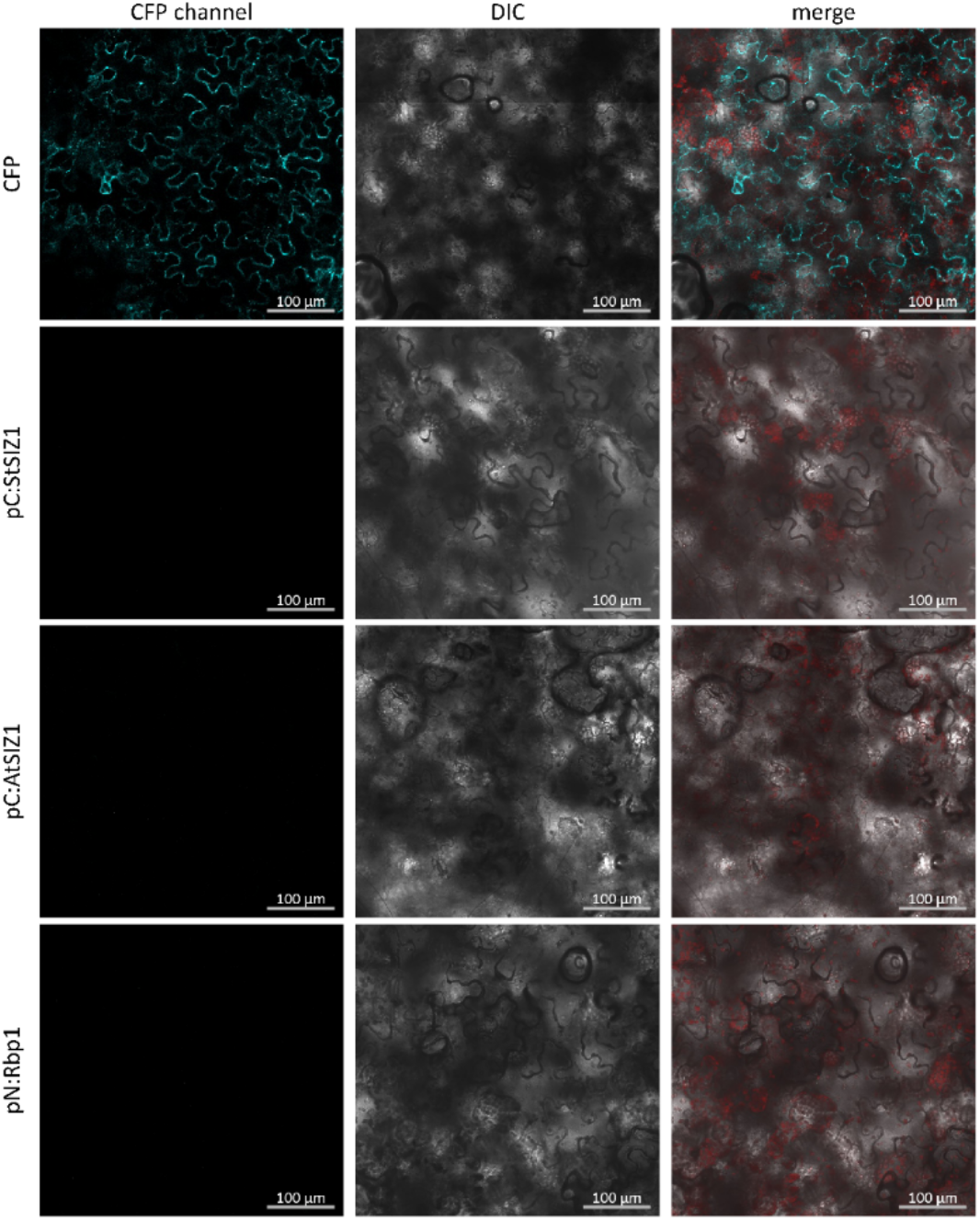
Individual bimolecular fluorescence complementation constructs do not emit fluorescence when infiltrated individually. Individual N-SCFP3 (pN) or C-SCFP3 (pC) fusions to StSIZ1, GpRbp-1, and. A CFP transformation is shown for comparison of the confocal microscopy settings.

**Supplemental Figure 4.**
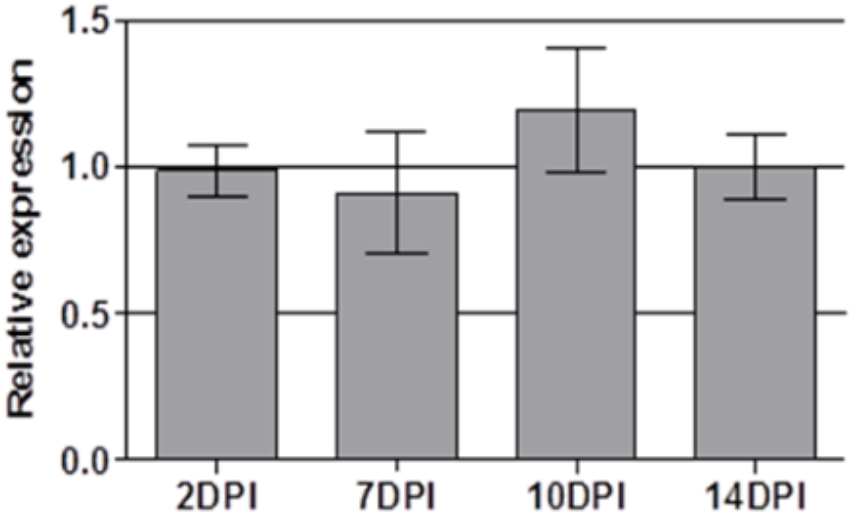
SIZ1 gene expression during nematode infection of Arabidopsis roots. AtSIZ1 expression as quantified by RT-PCR. Relative expression of AtSIZ1 is calculated relative to the geometric mean of reference genes UBP22 (Hofmann & Grundler, 2007) and UBQ5 (Anwer *et al*., 2018). Error bars indicate the standard error of 3 independent biological replicates.

**Supplemental Figure 5.**
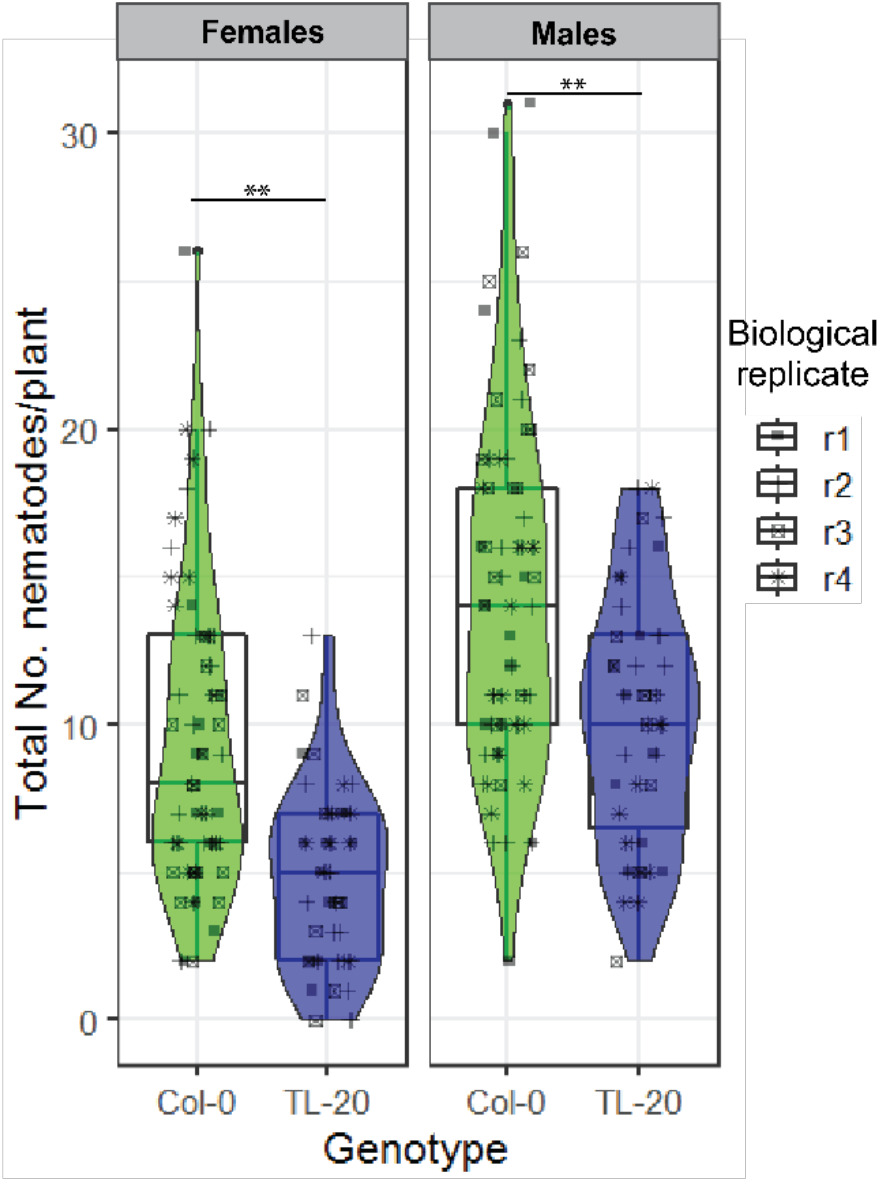
Less *H. schachtii* males and females infect the roots of *SIZ1-2 Arabidopsis.* Total amount of females and male nematodes present in the roots of *siz1-2* and wild-type (Columbia-0) *Arabidopsis,* after 2 weeks of infection. Whiskers indicate the maximum and minimum data points and violin plots describe the distribution of all data points. Results are combined measurements from 4 independent biological repeats. n_Col-0_= 65 and n_siz1-2_=47. Stars indicate statistical significance of the differences in the amount or size of nematodes infecting the roots of *siz1* line and the wild-type control, established by one-way ANOVA, (**p-value<0.001).

**Supplemental Table 2.** HiSat mapping of RNAseq reads to Arabidopsis genome.

| *Number* | *Aligned reads (%)* | *Multiple aligned reads (%)* | *Non-aligned reads (%)* | *Sample name* |
| --- | --- | --- | --- | --- |
| 0 | 92.76 | 3.89 | 3.35 | 1v2 |
| 1 | 87.83 | 3.86 | 8.3 | 2v2 |
| 2 | 93.33 | 3.77 | 2.9 | 3v2 |
| 3 | 83.68 | 3.4 | 12.92 | 4v2 |
| 4 | 89.92 | 5.13 | 4.95 | 5v2 |
| 5 | 82.08 | 3.28 | 14.64 | 6v2 |
| 6 | 92.32 | 3.95 | 3.73 | 7v2 |
| 7 | 86.23 | 3.22 | 10.55 | 8v2 |
| 8 | 90.44 | 5.52 | 4.03 | 9v2 |
| 9 | 93.66 | 3.63 | 2.71 | 11v2 |
| 10 | 85.71 | 3.29 | 11 | 12v2 |
| 11 | 87.56 | 4.23 | 8.21 | 10v2 |

**Supplemental Table 3. A list of all the genes significantly regulated in *siz1-2* plants**. genelD Indicates the TAIR ID for each entry; the test column indicates which comparison was made; the significance column gives the significance of the difference as determined by the linear model (with FDR correction), the effect column shows the size of the difference in gene expression (log2-units; negative values are lower expressed in col-0, positive values higher expressed in col-0). The significance_FDR column lists the q-values as determined by Benjamini-Hochberg correction. The columns thereafter list properties of the genes.

Available at; https://drive.google.com/open?id=1YktRjxXNRkLoFI3ZKyjasnUZU-bZ9g28

**Supplemental Table 4.**
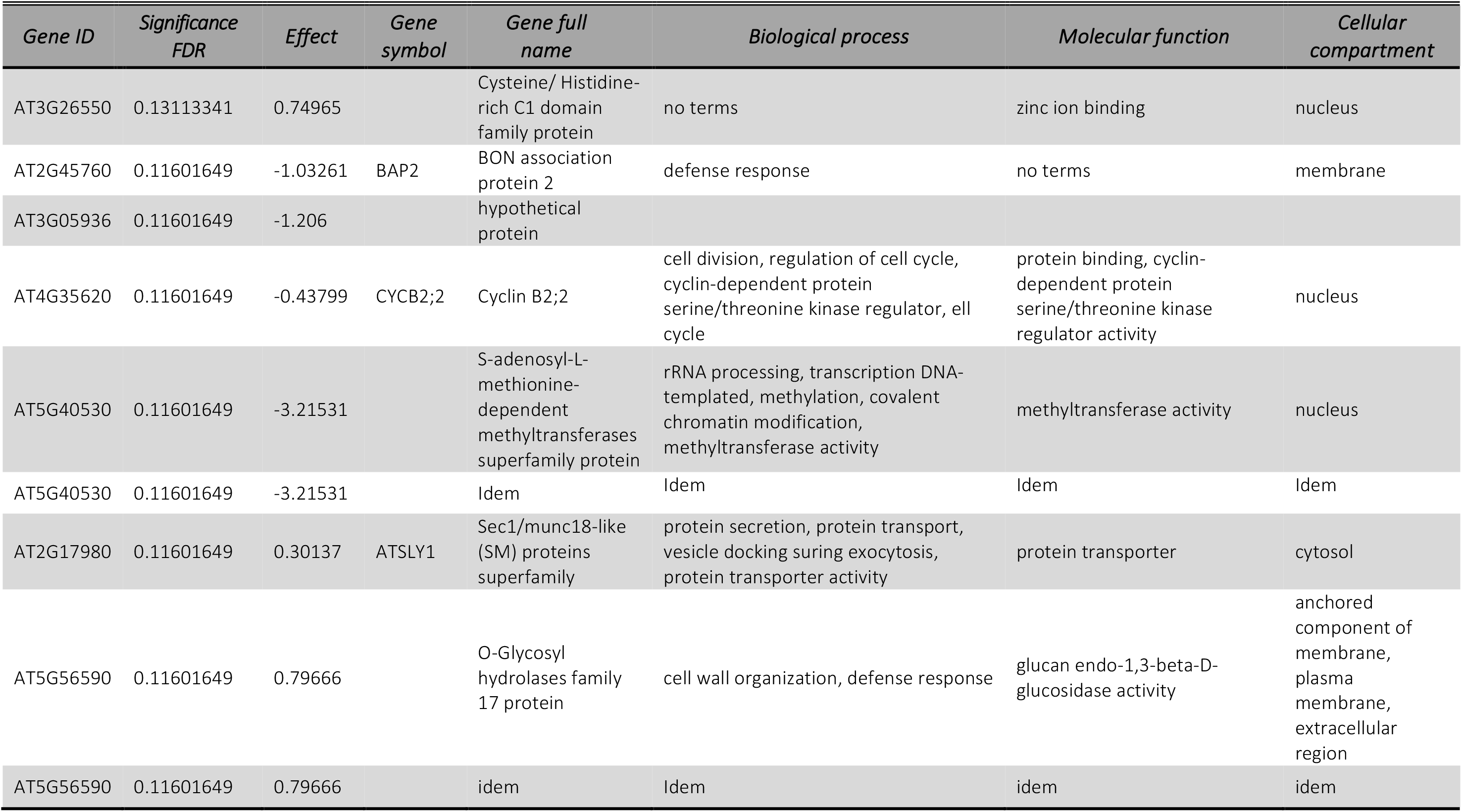
Genes significantly regulated by nematode infection and the mutation in *siz1-2* (interaction). The gene, significance FDR and effect are obtained from Supplemental Table 2. The Gene Symbol, Gene full-name and associated GO annotations were obtained from ThaleMine (Krishnakumar *et al*., 2014).

**Supplemental Table 5.**
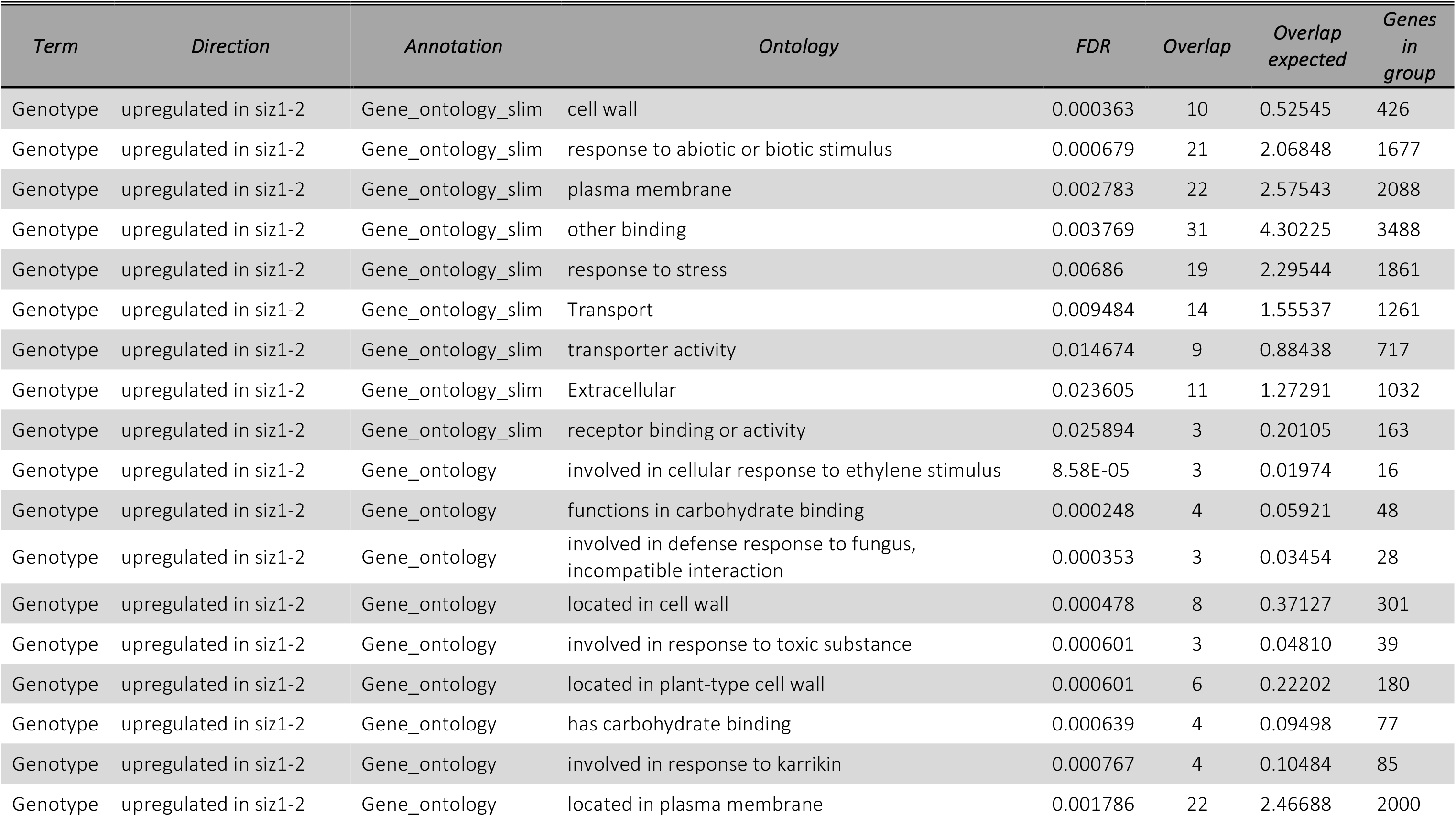
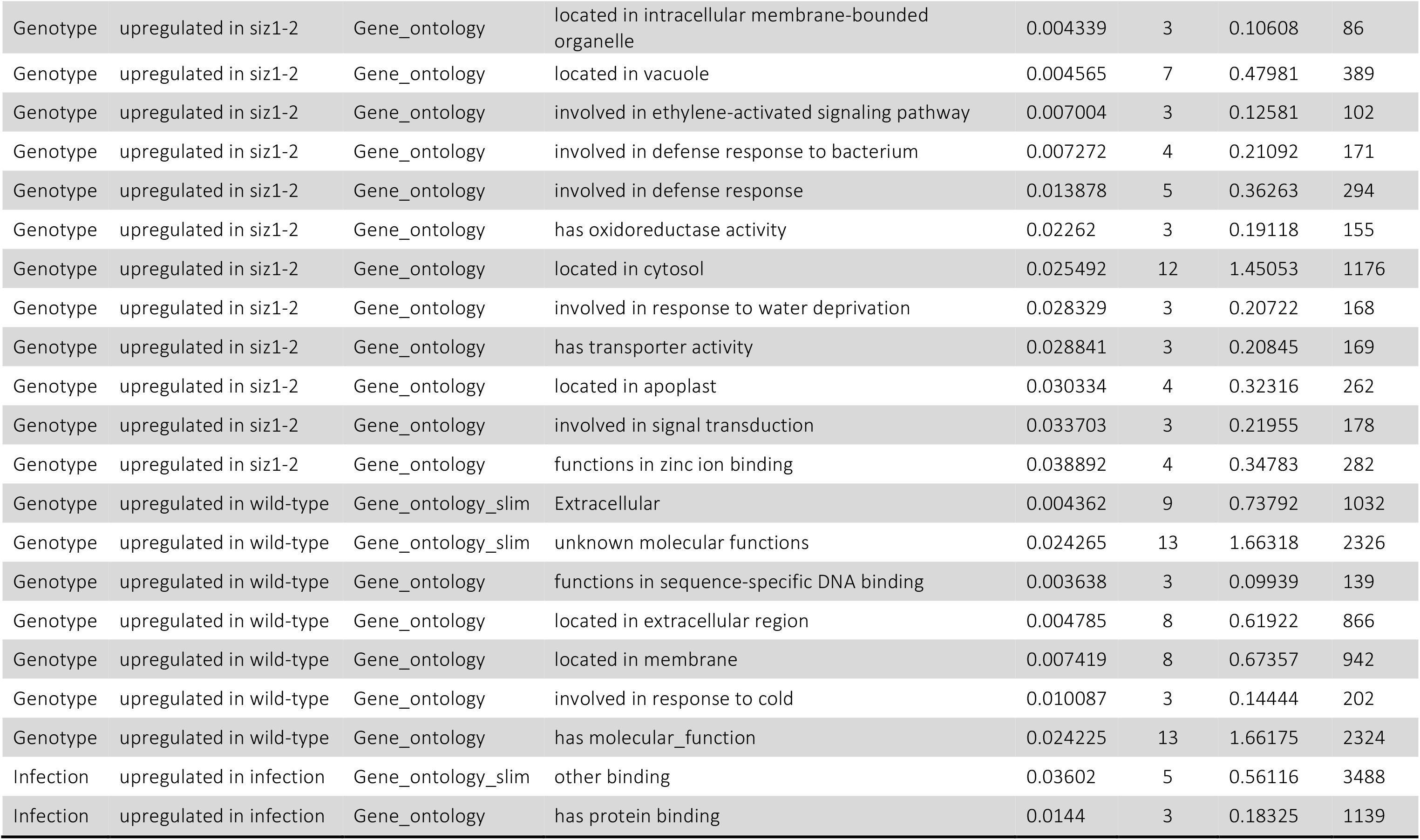
Gene Ontology categories enriched in genes differentially regulated either by the mutation in *SIZ1-2 Arabidopsis* or by infection with *H. schachtii.*

**Supplemental Figure 6.**
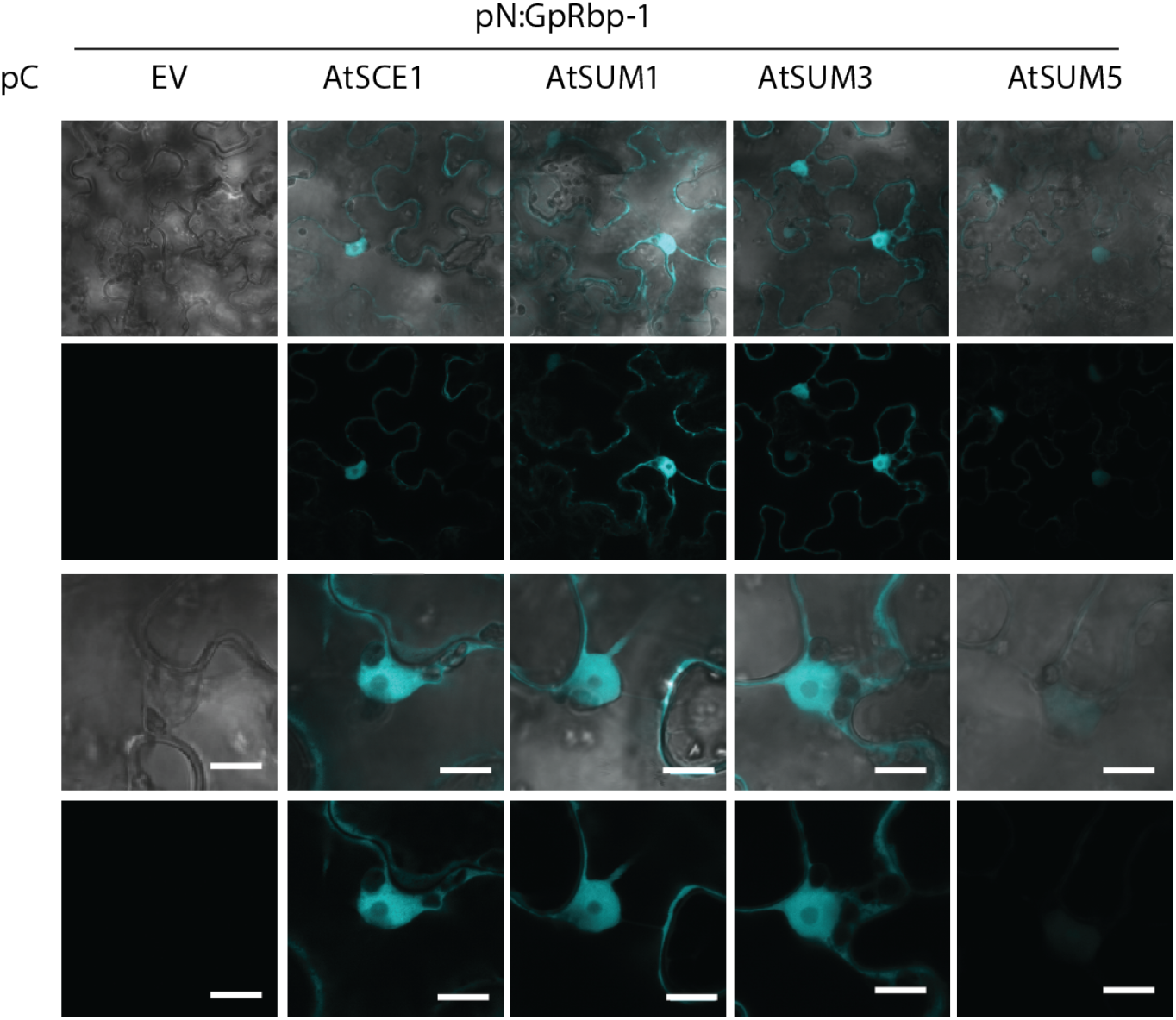
Gp2Rbp1 interacts with other members of the SUMO machinery. Bimolecular fluorescence complementation of N and C-terminal regions of SCFP3a. SCFP3A amino acids 1-173 were fused to GbRpb-1 (pN:Gp-Rbp1) and SCFP3A amino acids 156-239 were fused to SUMO1,3,5 or SCE1 (e.g. pC:AtSUM1). The corresponding pN/pC pairs were co-infiltrated to *N. benthamiana* leaves. Co-expression of pN:EV or pC:EV were used as negative controls. The CFP emission channel is shown in blue, light emission in white in the differential interference contrast channel and chloroplast auto-fluorescence is shown in red in the merge channel. Results are representative of 2 biological repeats and all co-infiltrations contain the silencing suppressor p19.

**Supplemental Figure 7.**
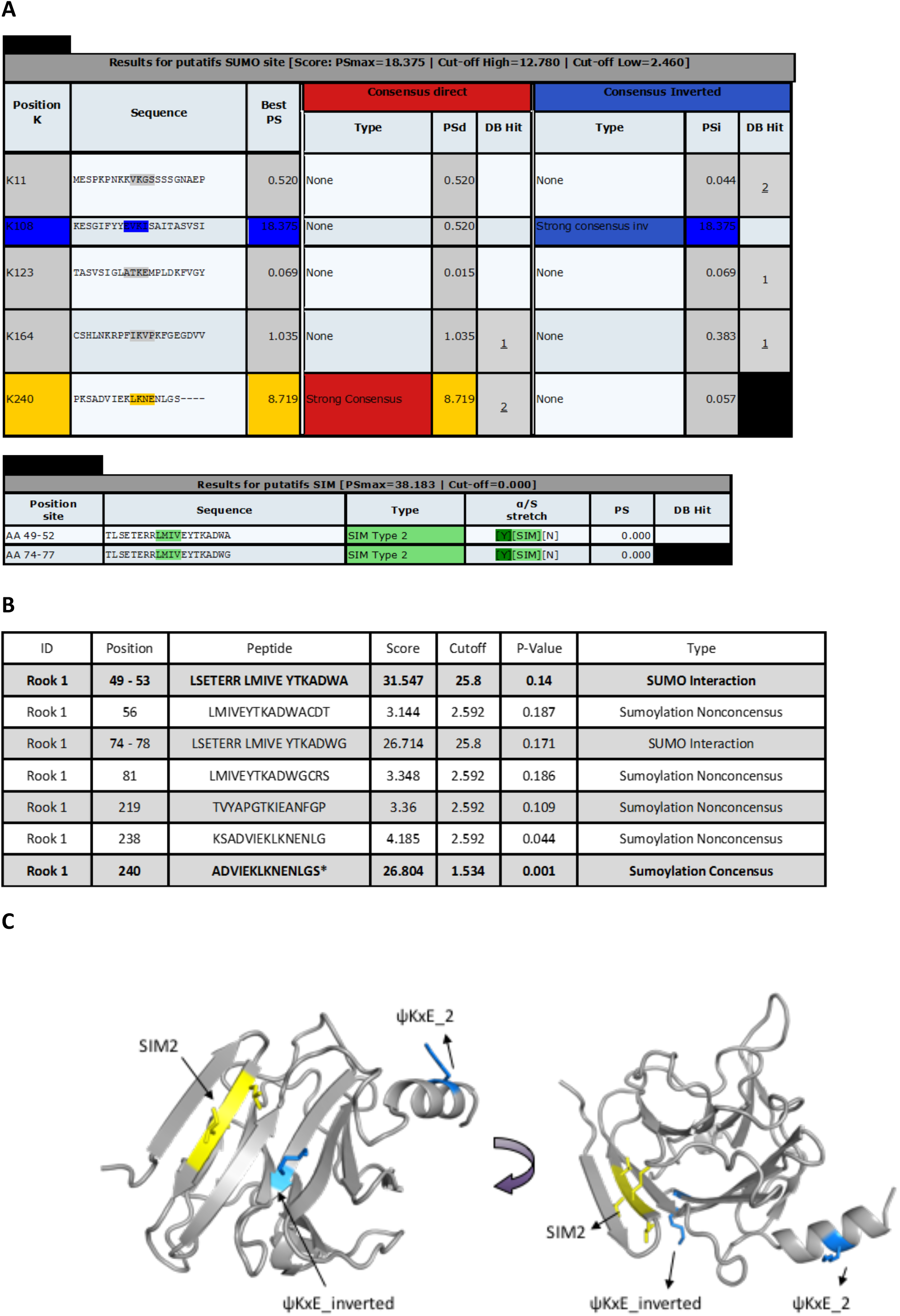
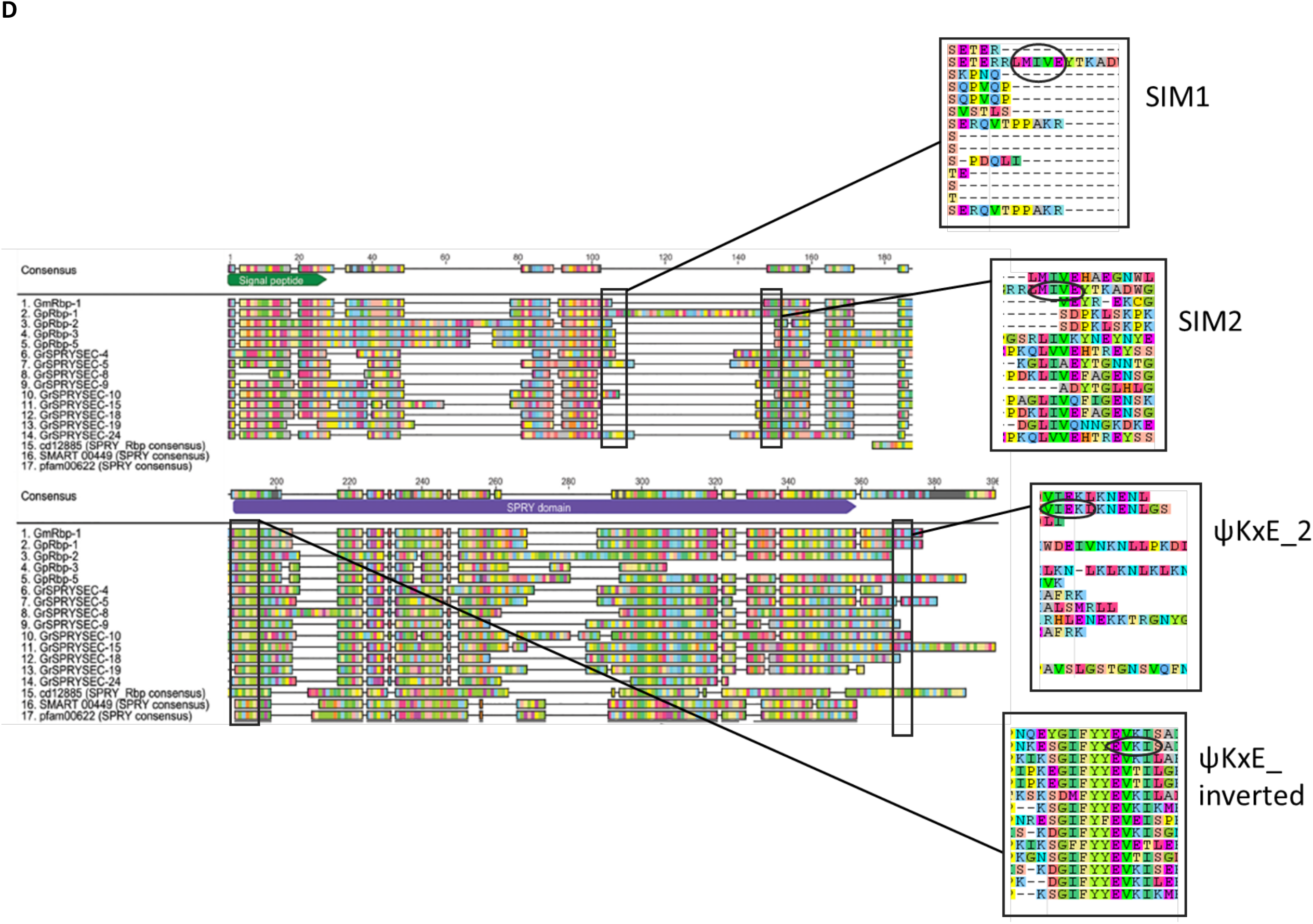
SUMOylation and SUMO interacting motifs are predicted in GpRbp-1. Lysine residues 108 and 240 of GpRbp-1 are predicted as SUMOylation sites. The amino acid stretch from position 49 to 53 and from position 74 to 78 of GpRbp-1 are predicted so function as SIMs, using **A)** The Joined Advanced SUMOylation site and SIM Analyser (JASSA) (Beauclair *et al*., 2015) and **B)** The GPS-SUMO Webserver tool (Zhao *et al*., 2014). **C)** Computational structural model of GpRbp-1 (grey), with SIM2 (yellow), and ΨKxE sites indicated (blue). GpRbp-1 was modelled by remote homology modelling from human RBPM10 (PDB: 5JI7) & SPRYD3 (PDB 2YYO) and from mouse RBP9 (PDB: 5J1A). **D.)** Location of SUMO and SIM sites in SPRYSEC alignment (modified from Diaz Granados, *et al,* 2016).

**Supplemental Table 6.**
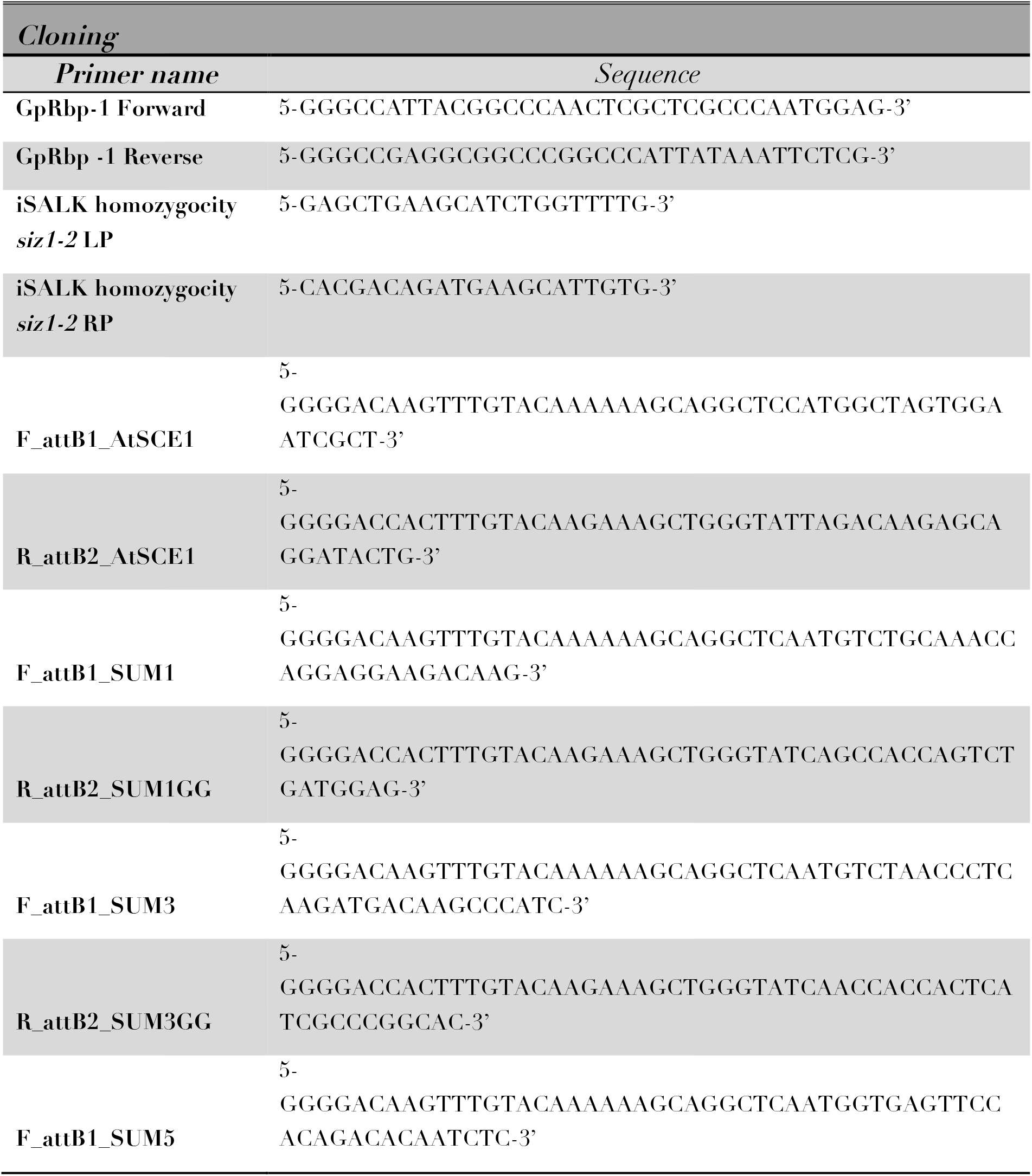
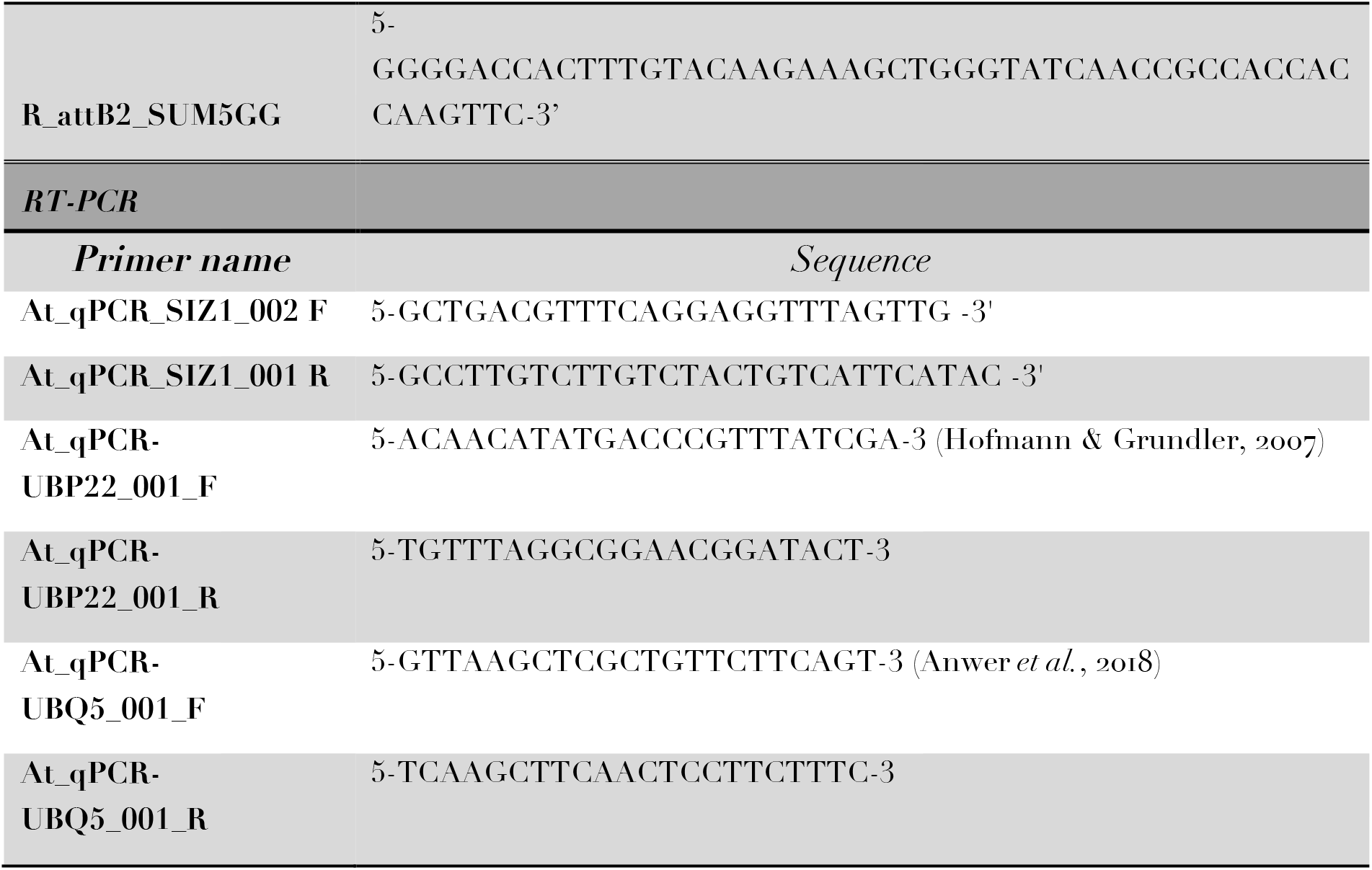
Primers mentioned in the text

